# Sidetrak: a de novo-designed miniprotein targets the kinesin-1 coiled-coil stalk to inhibit MAP7 binding and disrupt mitochondrial transport

**DOI:** 10.64898/2026.11.16.767073

**Authors:** Jessica A. Cross, Sarah A. Bristow, Thilani Babuji, Derek N. Woolfson, Mark P. Dodding

## Abstract

The coiled-coil domains of cytoskeletal motors contain regulatory surfaces that offer largely unexploited targets for manipulating intracellular transport. The microtubule-associated protein MAP7 promotes kinesin-1 motor activity, binding a coiled-coil surface that overlaps the TPR docking site (TDS), a regulatory interface involved in autoinhibition. Starting from an AlphaFold3 model of this relatively feature-poor surface, we used BoltzGen to design Sidetrak, a 36-amino acid miniprotein that binds kinesin-1 with nanomolar affinity. Binding requires the TDS and is abolished by mutation of the predicted Sidetrak interface, supporting the modelled local geometry of the targeted coiled-coil region. Sidetrak inhibits MAP7 binding, reduces MAP7-dependent kinesin-1 recruitment to microtubules in cells and causes perinuclear redistribution of mitochondria, a kinesin-1 cargo. Sidetrak binds all three kinesin-1 heavy-chain paralogues, and quantitative interaction proteomics revealed selective association with endogenous kinesin-1 complexes and associated cargo adaptors. These findings demonstrate selective disruption of a motor-MAP interaction by a de novo-designed protein and highlight regulatory coiled-coil surfaces as targets for manipulating intracellular transport. Beyond providing a means to manipulate intracellular transport, this work illustrates the potential of de novo protein design to test structural models of protein interfaces and probe endogenous protein networks to uncover underlying cellular mechanisms.

## Introduction

Intracellular transport by cytoskeletal motor proteins is tightly regulated to ensure the spatially and temporally controlled organisation of cellular components (*1*). Therefore, the ability to selectively manipulate motor activity provides a means to interrogate transport mechanisms and potentially correct pathological transport defects arising from motor dysregulation. The therapeutic potential of targeting molecular motors is exemplified by the development of small-molecule modulators of cardiac myosin II for the treatment of cardiomyopathies (*2*). These advances demonstrate that motor proteins are amenable to pharmacological modulation, although strategies targeting their ATPase motor domains have yet to achieve comparable success for motors mediating intracellular transport. Motor function, however, is also governed by diverse regulatory interactions outside the enzymatic core, including those controlling motor activation, cargo recognition and recruitment to cytoskeletal tracks. Recent advances in *de novo* protein design may offer opportunities to target these regulatory interfaces with high precision (*3–7*).

Kinesin-1, the prototypic kinesin family microtubule motor, comprises three closely related heavy-chain (KHC) paralogues, KIF5A, KIF5B and KIF5C, which associate with kinesin light chains (KLC1–4) to form heterotetrameric motor complexes. Kinesin-1 is regulated through conformational rearrangements of its extended coiled-coil scaffold that couple relief of autoinhibition to cargo recognition and microtubule engagement (*1, 5, 8–12*). We recently identified a regulatory site within the first KHC coiled coil (CC1), termed the TPR docking site (TDS), that contributes to both autoinhibition and motor activation (*12*). Structural modelling using AlphaFold3 (AF3) together with biochemical analysis, hydrogen–deuterium exchange mass spectrometry and negative-stain EM supports engagement of this site by the KLC TPR domains in the autoinhibited state. The TDS also provides a binding site for microtubule-associated protein 7 (MAP7), which promotes kinesin-1 recruitment and activation (*13–19*) (**Fig. 1A**). Therefore, this region represents a complex regulatory interface through which distinct protein interactions control kinesin-1 activity. Understanding how these interactions are coordinated, and how their effects on motor function can be selectively manipulated, presents an opportunity to dissect the mechanisms underlying kinesin-1 regulation.

**Figure 1.**
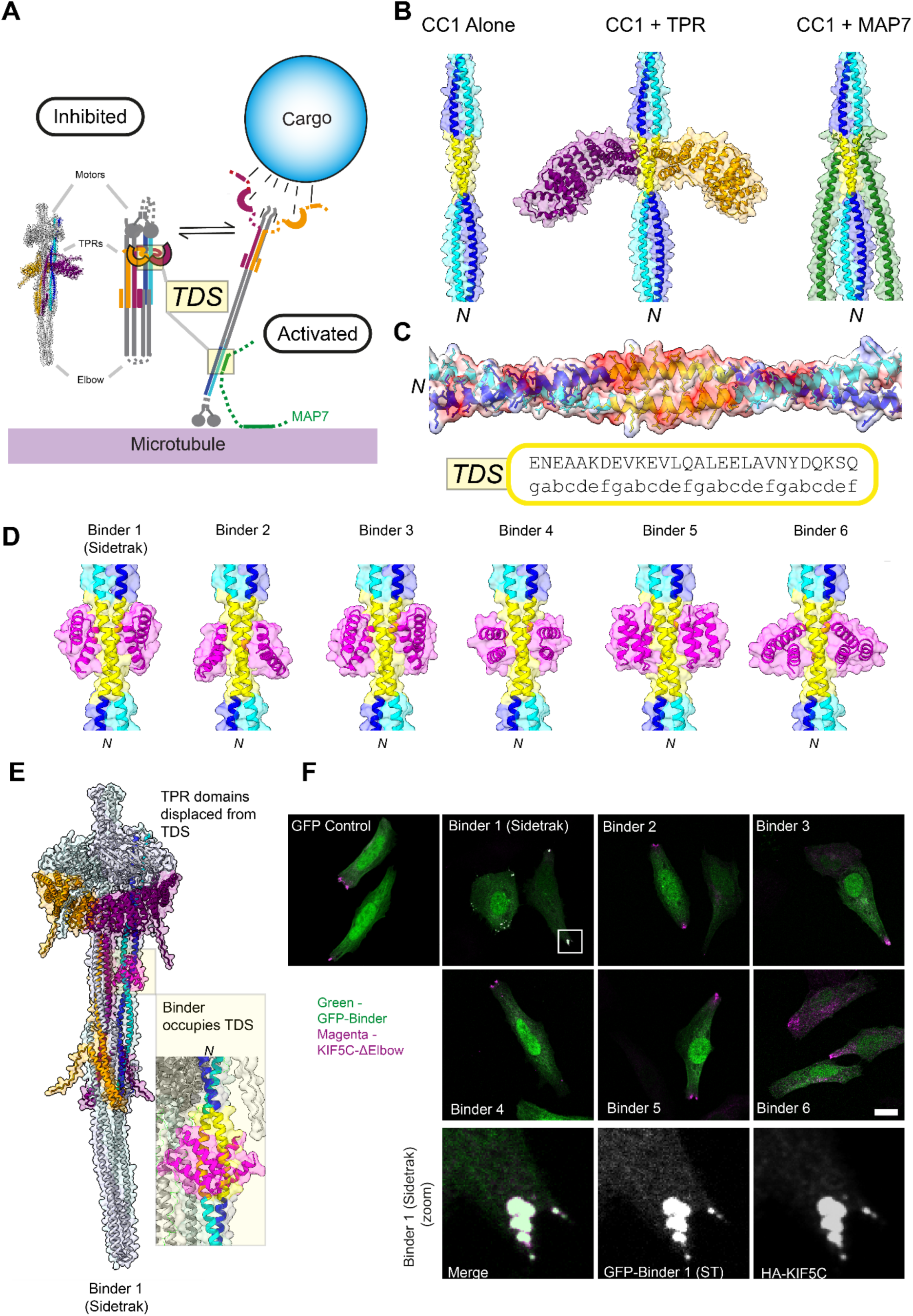
*De novo* design of a binder targeting the TPR docking site (TDS) on kinesin-1 CC1. **(A)** Schematic showing a generalised regulatory framework for kinesin-1 activation and the dual role of the TDS. In the absence of cargo, kinesin-1 adopts an inhibited conformation (modelled in AF3 and described in detail in (*12*)) in which the KLC TPR domains dock onto the TDS within KHC CC1. Cargo engagement through a variety of sites destabilises the inhibited state, cooperating in recruitment and activation with the microtubule-associated protein MAP7, which also binds the TDS. **(B)** AF3 models of KIF5C CC1 alone (NP_001101200.1; a.a. 410–558), in complex with the KLC TPR domains (NP_032476.2; a.a. 205–495), or with the kinesin-binding domain of MAP7 (NP_001185564.1; a.a. 467–610). The TDS, defined as four heptads of CC1 encompassing the TPR- and MAP7-binding sites, is shown in yellow (a.a. 475 to 502 of KIF5C; NP_001101200.1) **(C)** Electrostatic surface representation of CC1, together with its sequence and structure-based assignment of the coiled-coil heptad register from SOCKET2 (*54*). Surface is coloured by Coulombic electrostatic potential calculated in UCSF Chimera X. **(D)** AF3 models of six candidate binders modelled with CC1 as 2:2 heterotetramers. **(E)** AF3 model of Binder 1 (Sidetrak) bound to the full-length kinesin-1 heterotetramer. **(F)** Fluorescence images of HeLa cells co-expressing HA-KIF5C-ΔElbow (magenta) and the indicated GFP-tagged binder candidates. Scale bar, 10 µm. Enlarged panels below show colocalisation of Binder 1 (Sidetrak) with HA-KIF5C-ΔElbow at the cell periphery.

Despite this evidence for TPR engagement of the TDS, a recent cryo-EM study proposed a substantially different arrangement of the CC1 helices and alternative positions for the TPR domains, challenging this model of TPR engagement (*20*) **(Fig. S1).** The relatively low resolution of these data, obtained from chemically crosslinked kinesin-1 complexes, limits the extent to which the overall architecture and the conformation of this regulatory region can be unambiguously defined. Moreover, given the dynamic and flexible nature of kinesin-1, chemical crosslinking could stabilise a non-native arrangement of its regulatory domains. In the AF3 model, the TDS forms a canonical homodimeric coiled coil, presenting a relatively feature-poor surface for molecular recognition. This model provides an *a priori* structural hypothesis that can be tested through binder design: i.e. the successful generation of a binder that engages the predicted interface would support the modelled local geometry. The TDS offers an opportunity to use *de novo* protein design to both probe and manipulate a regulatory interface for which high-resolution structural information is unavailable.

Here, we use AlphaFold3 (*21*) and BoltzGen (*22*) to design a miniprotein targeting the TDS. The resulting binder, *Sidetrak,* binds kinesin-1 with high affinity and specificity through the predicted interface and inhibits its interaction with MAP7. Sidetrak engages all three KIF5 paralogues and enables the recovery of endogenous kinesin-1 complexes and associated proteins for proteomic analysis. Functionally, Sidetrak disrupts MAP7-dependent recruitment of kinesin-1 to microtubules and alters the distribution of both kinesin-1 and mitochondria, a kinesin-1 cargo. These findings demonstrate that protein design can manipulate endogenous motor function through a regulatory interface outside the conserved motor domain while providing experimental support for the predicted local architecture of CC1. More broadly, they highlight motor–MAP interactions and coiled-coil surfaces in motor proteins as tractable targets for synthetic modulation.

## Results

### Ligand-constrained modelling and cellular screening identify a *de novo* kinesin-1 binder

To establish a structural template for binder design, we used AF3 to model interactions of the kinesin-1 CC1 TDS (KIF5C) with TPR domains from KLC1 and MAP7, reasoning that these natural binding partners could help define the local structure of CC1. As previously reported, AF3 predicted that both the KLC TPR domains and the MAP7 kinesin-binding domain (KBD) engage overlapping surfaces within the TDS of the KHC CC1 homodimer (*12*) (**Fig. 1B, Fig. S2**). The TPR domain engaged CC1 through its first α-helical hairpin, oriented approximately perpendicular to the CC1 axis, whereas MAP7 bound as a single α-helix oriented approximately parallel to CC1. The local CC1 structure was modelled with greater confidence and more consistent side-chain positions in the presence of either binding partner than in models of CC1 alone, providing a partner-constrained conformation for binder design.

We used the CC1 conformation from the AF3 CC1–TPR complex as the design template. The TPR domains were computationally removed to expose the TDS, an overall negatively charged binding surface (**Fig. 1C**). Using BoltzGen, we generated 10,000 candidate binders of 30–50 amino acids targeted to this surface, of which six passed default filters and were selected for further analysis (**Fig. S3**). AF3 modelling with the CC1 homodimer predicted that binders 1–4 formed short α-helical hairpins, whereas binders 5 and 6 formed three-helix bundles, with all six engaging the intended TDS (**Fig. 1D, Fig. S4**). Despite differences in geometry, the designs recapitulated features of the TPR–TDS interface, including similar distributions of charge and hydrophobicity (**Fig. S5**). Modelling in the context of the complete kinesin-1 heterotetramer further predicted that all six binders engaged the TDS rather than alternative sites within the complex, displacing the KLC TPR domains from this surface (**Fig. 1E, Fig. S6**).

To screen the six candidate TDS binders experimentally, we expressed each as an N-terminal GFP fusion in HeLa cells together with HA-tagged KIF5C-ΔElbow, a previously described constitutively active kinesin-1 mutant in which disruption of a flexible “elbow” region relieves autoinhibition and promotes motor activity (*10*). This mutant accumulates at the cell periphery, providing a cellular readout for the recruitment of interacting proteins. We reasoned that binders interacting robustly with KIF5C-ΔElbow would be recruited to these peripheral accumulations. Of the six designs, binder 1, the top-ranked candidate, showed strong recruitment, whereas GFP alone and the remaining designs showed little or no detectable enrichment above background (**Fig. 1F)**. Therefore, we prioritised binder 1, hereafter termed Sidetrak, for further characterisation.

### Sidetrak binds the kinesin-1 TDS with high affinity

To determine whether Sidetrak binds kinesin-1 directly, we measured binding of TAMRA-labelled Sidetrak peptide to K560, a truncated kinesin-1 construct encompassing the motor domain and CC1, including the TDS. Fluorescence polarisation revealed an apparent dissociation constant (*K*d) of 110 ± 13 nM (**Fig. 2A, Fig. S7**), indicating more than an order of magnitude tighter binding than previously reported for MAP7 or KLC TPR domains, whose reported affinities range from low to high micromolar depending on the paralogues and experimental conditions (*12, 13*). No binding was detected to K401, a shorter construct that lacks the TDS (**Fig. S7**), supporting a requirement for this region in Sidetrak binding. In cells, GFP-Sidetrak co-precipitated both wild-type HA-KIF5C and the constitutively open HA-KIF5C-ΔElbow mutant, each in complex with HA-KLC2 (**Fig. 2B**). Sidetrak showed a modest, approximately two-fold preference for KIF5C-ΔElbow. This preference was the same in direction, but smaller in magnitude, to than previously observed for MAP7 and is consistent with recognition of a region of CC1 whose accessibility is regulated by the conformational state of kinesin-1 (*12*).

**Figure 2.**
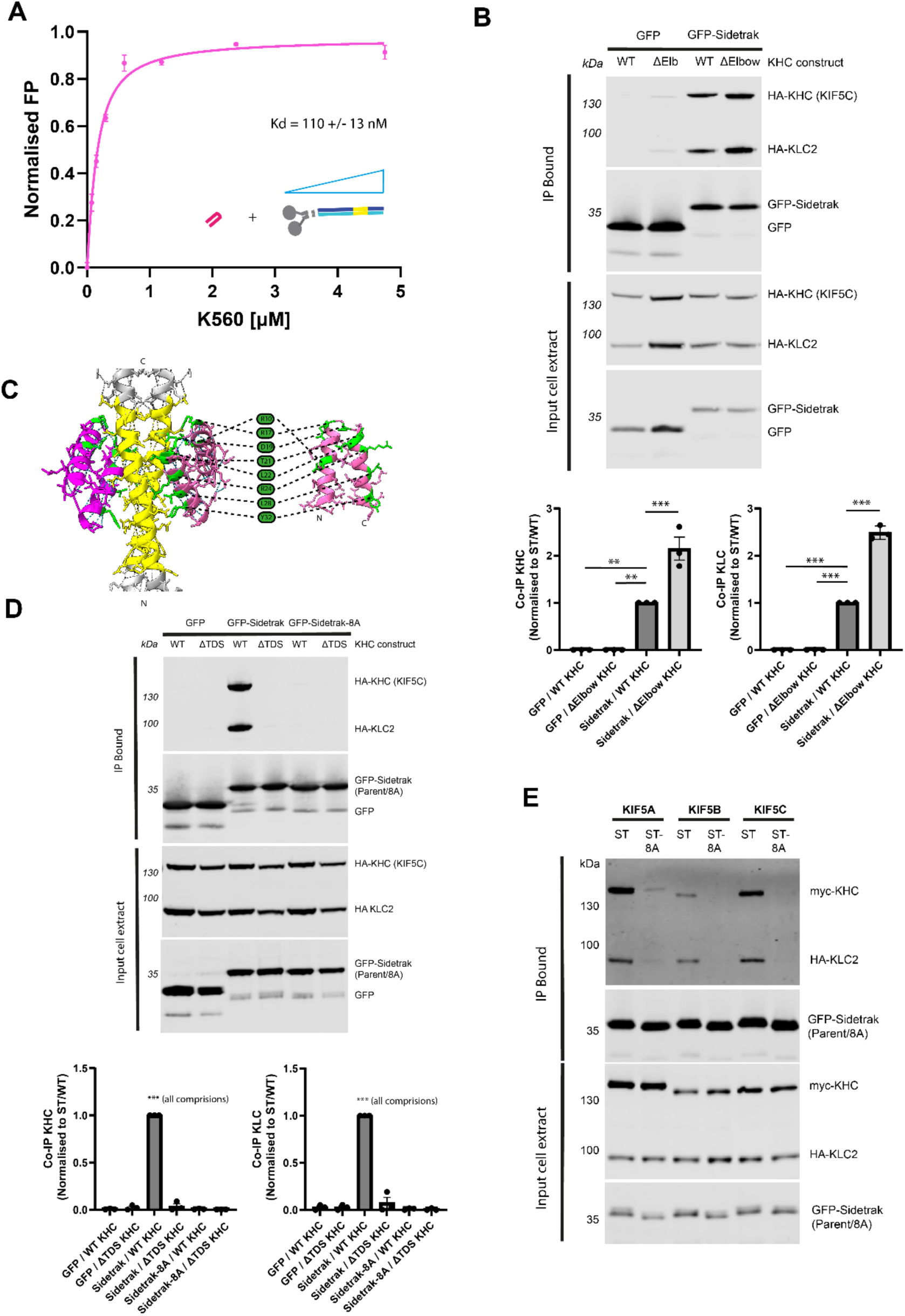
Sidetrak directly binds KHC and targets the TDS. (A) Fluorescence polarisation analysis of TAMRA-labelled Sidetrak peptide binding to recombinant K560, encompassing the KHC motor domain and proximal coiled-coil region, including the TDS. (B) Co-immunoprecipitation analysis of GFP-Sidetrak binding to wild-type HA-KIF5C/HA-KLC2 or the constitutively open HA-KIF5C-ΔElbow/HA-KLC2 mutant. Sidetrak bound both constructs, with increased binding to KIF5C-ΔElbow. (C) AF3 model of Sidetrak (magenta) bound to the KIF5C CC1 TDS (yellow), highlighting Sidetrak residues predicted to contribute to the binding interface using BUDE computational alanine scanning of Sidetrak and mutated to alanine in Sidetrak-8A. (D) Reciprocal perturbation of the predicted Sidetrak–TDS interface. GFP-Sidetrak binding was assessed using wild-type KIF5C or KIF5C-ΔTDS, in which the TDS is replaced by four heptads of the *de novo*-designed coiled coil CC-Di, together with wild-type Sidetrak or the Sidetrak-8A interface mutant. Disruption of either the KHC TDS or the predicted Sidetrak-binding interface disrupted the interaction. (E) Co-immunoprecipitation analysis of GFP-Sidetrak binding to myc-tagged KIF5A, KIF5B and KIF5C co-expressed with HA-KLC2. Wild-type Sidetrak bound all three KHC paralogues, whereas binding of the Sidetrak-8A interface mutant was disrupted. Data are presented as mean ± SEM from *n* = 3 independent experiments, with individual experiments shown. Statistical significance was determined by one-way ANOVA followed by Dunnett’s multiple-comparisons test, comparing each condition with the indicated WT control. **P < 0.01, ***P < 0.001.

We tested the predicted Sidetrak-TDS interface by independently disrupting the binding surface on both the target and binder. For the target, we used the previously characterised KIF5C-ΔTDS construct, in which the TDS is replaced by four heptads of the *de novo*-designed coiled coil cc-Di (*12*). This mutation was designed to preserve the predicted coiled-coil architecture while removing the native TDS side chains required for molecular recognition. For the binder, BUDE computational alanine scanning (*23, 24*) identified eight Sidetrak residues predicted to contribute substantially to CC1 binding (ΔΔG ≥ 5 kJ/mol), spanning both helices and contacting both chains of CC1 (**Fig. 2C, Fig. S8, Fig. S9**). To disrupt these interactions, simultaneously we substituted these residues with alanine to generate Sidetrak-8A. The substitutions were confined to the predicted CC1-binding surface (**Fig. S8**). Both KIF5C-ΔTDS and Sidetrak-8A disrupted the interaction in co-immunoprecipitation experiments (**Fig. 2D**). Together, these results support the designed model in which Sidetrak engages the kinesin-1 TDS and provide Sidetrak-8A as a negative control for subsequent experiments.

Next, we asked whether Sidetrak binds all three mammalian kinesin-1 heavy-chain paralogues, KIF5A, KIF5B and KIF5C. The TDS is highly conserved across all three KHCs despite greater sequence divergence in the surrounding CC1 region (**Fig. S10A**), and AF3 predicted Sidetrak to engage each through an essentially identical binding mode (**Fig. S10B**). Consistent with these predictions, GFP-Sidetrak co-immunoprecipitated myc-tagged KIF5A, KIF5B and KIF5C co-expressed with HA-KLC2, whereas the Sidetrak-8A interface mutant disrupted binding to all three (**Fig. 2E**). Thus, Sidetrak is a pan-kinesin-1 binder.

### Sidetrak inhibits MAP7 binding and MAP7-mediated recruitment of kinesin-1 to microtubules

The predicted Sidetrak-binding site overlaps with the MAP7-binding site on CC1, suggesting that Sidetrak should compete with MAP7 for kinesin-1 engagement. To test this, increasing concentrations of Sidetrak peptide were added to lysates containing GFP-MAP7 and HA-tagged KHC/KLC complexes prior to GFP-MAP7 immunoprecipitation. Sidetrak caused a concentration-dependent reduction in kinesin-1 co-immunoprecipitation with MAP7, with half-maximal displacement at 86 nM (95% CI 62–116 nM) for KHC and 79 nM (95% CI 36– 150 nM) for KLC2, and a maximal reduction of approximately 85% (**Fig. 3A**). These half-maximal displacement concentrations are close to the apparent affinity of Sidetrak for K560 measured by fluorescence polarisation (110 ± 13 nM), consistent with Sidetrak competing with MAP7 for an overlapping binding site on CC1. As an orthogonal approach, we determined whether Sidetrak inhibited MAP7-dependent kinesin-1 recruitment to microtubules in cells. Exogenous mCherry-MAP7 strongly recruited HA-KHC to MAP7-positive microtubule bundles, consistent with previous reports (*13, 17*). This recruitment was significantly reduced by GFP-Sidetrak but preserved with the binding-defective GFP-Sidetrak-8A mutant (**Fig. 3B, 3C**). Together, these data show that Sidetrak engages the TDS and competitively inhibits MAP7 binding and MAP7-dependent recruitment of kinesin-1 to microtubules.

**Figure 3.**
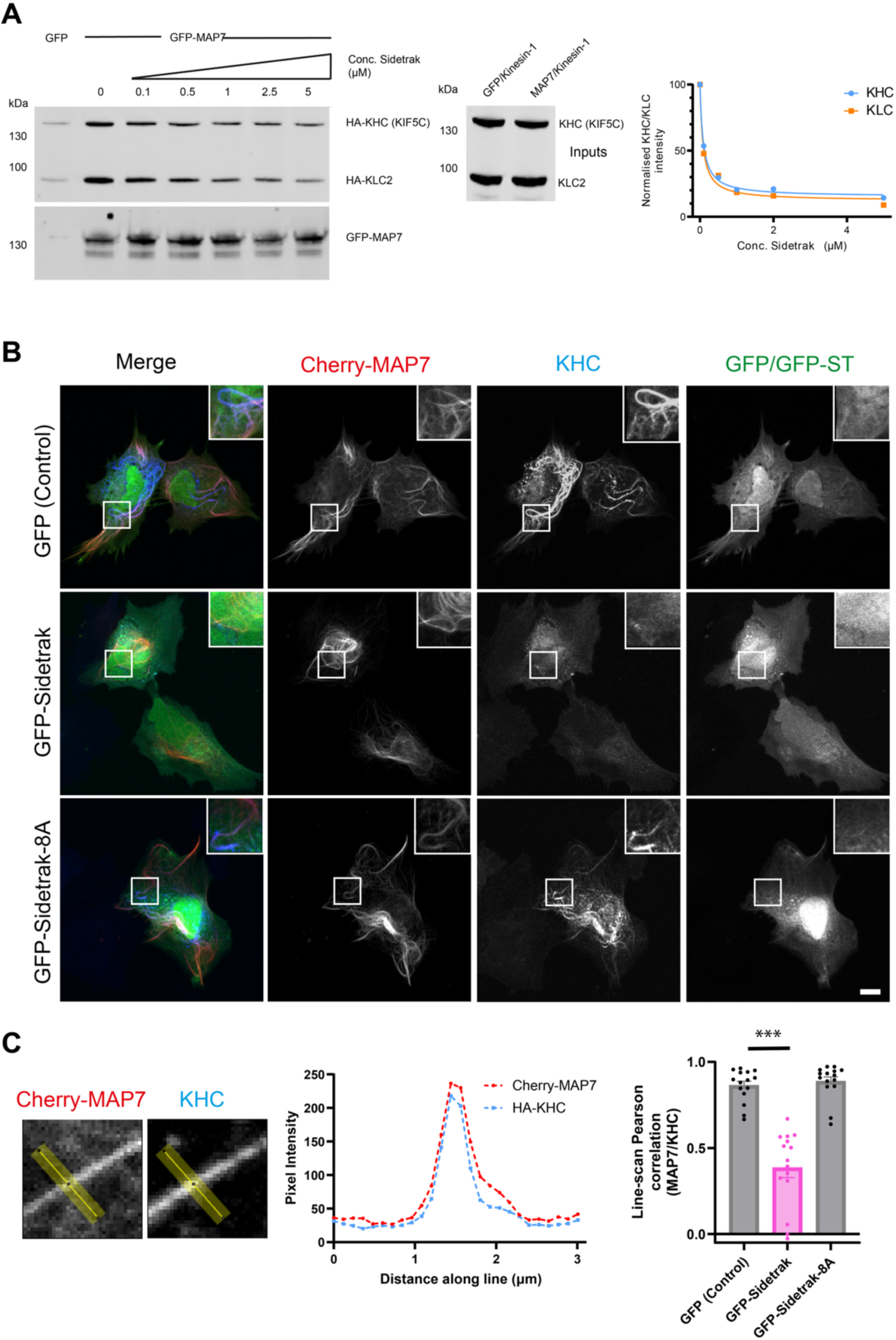
Sidetrak inhibits the interaction between MAP7 and KHC. (A) GFP-MAP7 and HA-tagged KHC/KLC complexes were co-expressed in cells. Increasing concentrations of Sidetrak peptide were added to cell lysates prior to immunoprecipitation of GFP-MAP7, and co-immunoprecipitated kinesin-1 was analysed by immunoblotting. Graph shows quantification of KHC and KLC2 co-immunoprecipitation relative to GFP-MAP7, normalised to the untreated control (100%) and fitted to a four-parameter inhibition model with the top constrained to 100%. IC50 values with profile-likelihood 95% confidence intervals are given in the text. Data are from one experiment representative of three independent experiments. (B) Representative fluorescence images of cells co-expressing mCherry-MAP7 and HA-KHC together with GFP, GFP-Sidetrak or the binding-defective GFP-Sidetrak-8A mutant. Exogenous MAP7 recruits KHC to MAP7-positive microtubules in control cells. This recruitment is inhibited by GFP-Sidetrak but retained in cells expressing GFP-Sidetrak-8A. (C) KHC recruitment to MAP7-positive microtubules was quantified by calculating the Pearson correlation between MAP7 and KHC fluorescence intensity profiles along 3-µm line scans (5-pixel width) drawn perpendicular to MAP7-positive microtubules. A representative (GFP-control) image and corresponding fluorescence intensity profiles are shown. Three microtubules were analysed per cell, and the resulting Pearson correlation coefficients were averaged to generate a single value per cell (n = 15 cells per condition). Data are mean ± SEM. Statistical significance was determined by one-way ANOVA with Dunnett’s multiple-comparisons test. \*\*\**P* < 0.001. Scale bars, 10 µm.

### Unbiased proteomic analysis reveals selective engagement of the kinesin-1 family and associated transport complexes

To assess the selectivity of Sidetrak, we performed quantitative comparative proteomic analysis of proteins co-immunoprecipitated with GFP-Sidetrak, GFP-Sidetrak-8A and GFP alone from HEK293 cells. Sidetrak strongly enriched kinesin-1 components relative to both negative controls, whereas Sidetrak-8A showed a profile similar to GFP alone (**Fig. 4A–G**). All three heavy-chain paralogues (KIF5A–C) and all four light chains (KLC1–4) were among the most prominent hits (Fig. 4A, B) and were recovered at high abundance (**Fig. 4F, G),** demonstrating engagement of endogenous kinesin-1 complexes. Apart from modest enrichment of KIF7, a non-motile kinesin-4 family member, no other cytoskeletal motors were significantly enriched.

**Figure 4.**
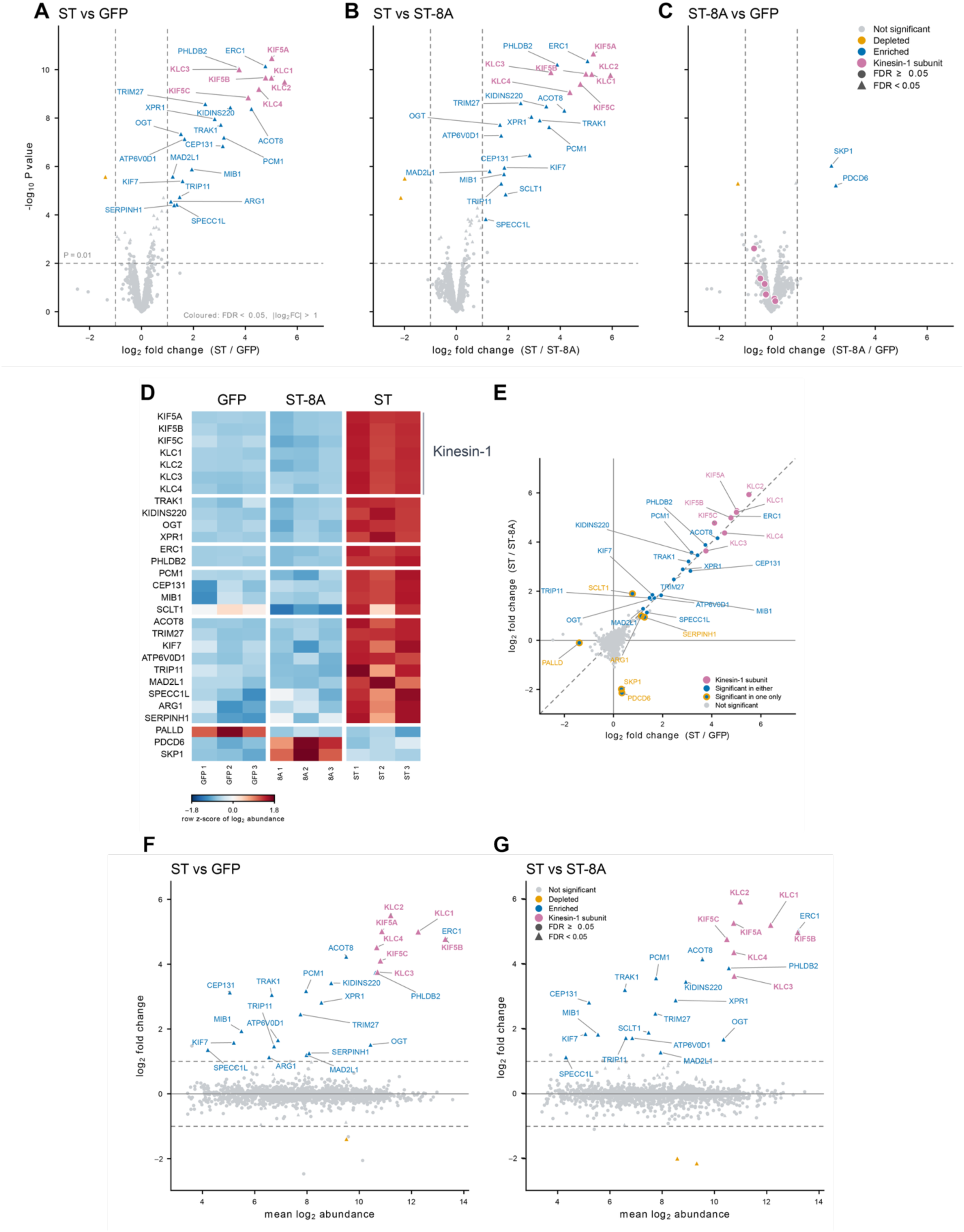
Quantitative proteomics identifies endogenous kinesin-1 complexes as the predominant targets selectively engaged by Sidetrak. **(A–C)** Proteins associated with GFP-Sidetrak (ST), binding-defective GFP-Sidetrak-8A (ST-8A) and GFP alone immuno-precipitated from HEK293 cells were analysed by tandem mass tag (TMT) quantitative proteomics, with three independent immunoprecipitates per condition. Protein abundances were log₂-transformed and median-normalised. Statistical comparisons used moderated t-tests from a single LIMMA model incorporating all three conditions, with Benjamini–Hochberg correction for multiple testing. Only protein groups supported by at least three unique peptides are shown. Volcano plots comparing ST with GFP (A), ST with ST-8A (B), and ST-8A with GFP (C). Axes show log₂ fold change and −log₁₀ unadjusted P value. Enriched and depleted proteins meeting FDR <0.05 and |log₂ fold change| >1 are coloured blue and orange, respectively; kinesin-1 subunits are highlighted in magenta. Triangles indicate FDR <0.05 and circles indicate FDR ≥0.05. Dashed lines mark log₂ fold changes of ±1 and P =0.01. **(D)** Heatmap showing row-wise Z-scores of log₂ protein abundances across individual immunoprecipitates. For each protein, values were centred on its mean across all nine samples and divided by its standard deviation. Colours therefore indicate relative abundance within each protein, rather than absolute abundance differences between proteins. **(E)** Comparison of log₂ fold changes for ST relative to GFP and ST-8A. Proteins meeting the enrichment criteria in both comparisons are shown in blue, those meeting the criteria in only one comparison in orange, and kinesin-1 subunits in magenta. The diagonal dashed line indicates equal fold changes in the two comparisons. **(F,G)** MA plots (M, log ratio; A, mean log signal) showing log₂ fold change against mean log₂ normalised TMT abundance for ST versus GFP (F) and ST versus ST-8A (G). TMT abundance represents relative protein-associated MS signal rather than absolute protein concentration. Colours and symbols are as in A–C; dashed lines mark log₂ fold changes of ±1.

Several additional proteins were consistent with recovery of kinesin-1-associated complexes. These included the kinesin-1 adaptors TRAK1 and KIDINS220, which bind directly to the motor complex (*25–30*), and were enriched approximately 9- and 11-fold, respectively, relative to Sidetrak-8A. Their respective binding partners OGT and XPR1 were also recovered (*31–33*). ERC1 (ELKS) and PHLDB2 (LL5β) were also strongly enriched. Both participate in the cortical capture of kinesin-1-driven Rab6-positive carriers (*34, 35*) with ERC1 also associating with the carriers en route to the plasma membrane (*36*). Hits also included a group of centriolar satellite-associated proteins: the scaffold PCM1, CEP131 and the E3 ubiquitin ligase MIB1, consistent with recovery of satellite-associated complexes (*37*). Their recovery suggests an association with centrosome-related machinery, of interest given the established roles of kinesin-1 in centrosome positioning (*38, 39*).

ACOT8, a predominantly peroxisomal acyl-CoA thioesterase (*40*) was also enriched. Although ACOT8 has no established direct connection to kinesin-1, it was previously identified alongside KIDINS220 and XPR1 in the SNX27 interactome of T lymphocytes (*41*), raising the possibility that it is recovered through an indirect association with trafficking complexes. TRIM27, an E3 ubiquitin ligase that promotes WASH-dependent endosomal actin assembly and retromer-mediated trafficking, was also enriched, providing a further potential link to the endosomal sorting machinery involved in KIDINS220 recycling (*42*).

Together, these data support selective engagement of endogenous kinesin-1 complexes by Sidetrak. Although off-target associations cannot be excluded, the recovery of proteins with known links to kinesin-1 and intracellular trafficking suggests that Sidetrak may provide a tool for identifying components of endogenous kinesin-1 transport complexes.

### Sidetrak promotes perinuclear accumulation of mitochondria and kinesin-1

Having established that Sidetrak selectively engages endogenous kinesin-1 and inhibits MAP7 binding, next we asked whether it could perturb a cellular process regulated by kinesin-1 and MAP7. Mitochondrial distribution provides a functional readout in this context, as loss of either KIF5B or MAP7 family proteins causes mitochondria to redistribute from the cell periphery towards the perinuclear region (*13*). Furthermore, TRAK1, a mitochondrial cargo adaptor, was enriched in our proteomic analysis, providing an additional rationale for investigating mitochondrial transport. In HeLa cells, GFP-Sidetrak similarly caused pronounced perinuclear redistribution of mitochondria compared with GFP-expressing controls (**Fig. 5A**). Quantification of the cumulative radial distribution confirmed this inward shift, with a significantly greater fraction of mitochondrial fluorescence within the central 50% of the cell (**Fig. 5B, C**). In contrast, the binding-defective GFP-Sidetrak-8A variant did not alter mitochondrial distribution, indicating that the phenotype depends on the designed kinesin-1-binding interface. Similar perinuclear redistribution was observed in HEK293 and U2OS cells **(Fig. S11),** while gross microtubule architecture remained unaffected **(Fig. S12).** GFP-Sidetrak itself also accumulated prominently in the perinuclear region, unlike the controls. Co-staining for endogenous KIF5B and KIF5C revealed that kinesin-1 accumulated at these GFP-Sidetrak-positive foci, indicating that Sidetrak-associated redistribution of the motor accompanies the changes in mitochondrial distribution (**Fig. 5D, E**). Together, these data show that targeting the TDS with Sidetrak is sufficient to drive perinuclear redistribution of both mitochondria and endogenous kinesin-1 in cells.

**Figure 5.**
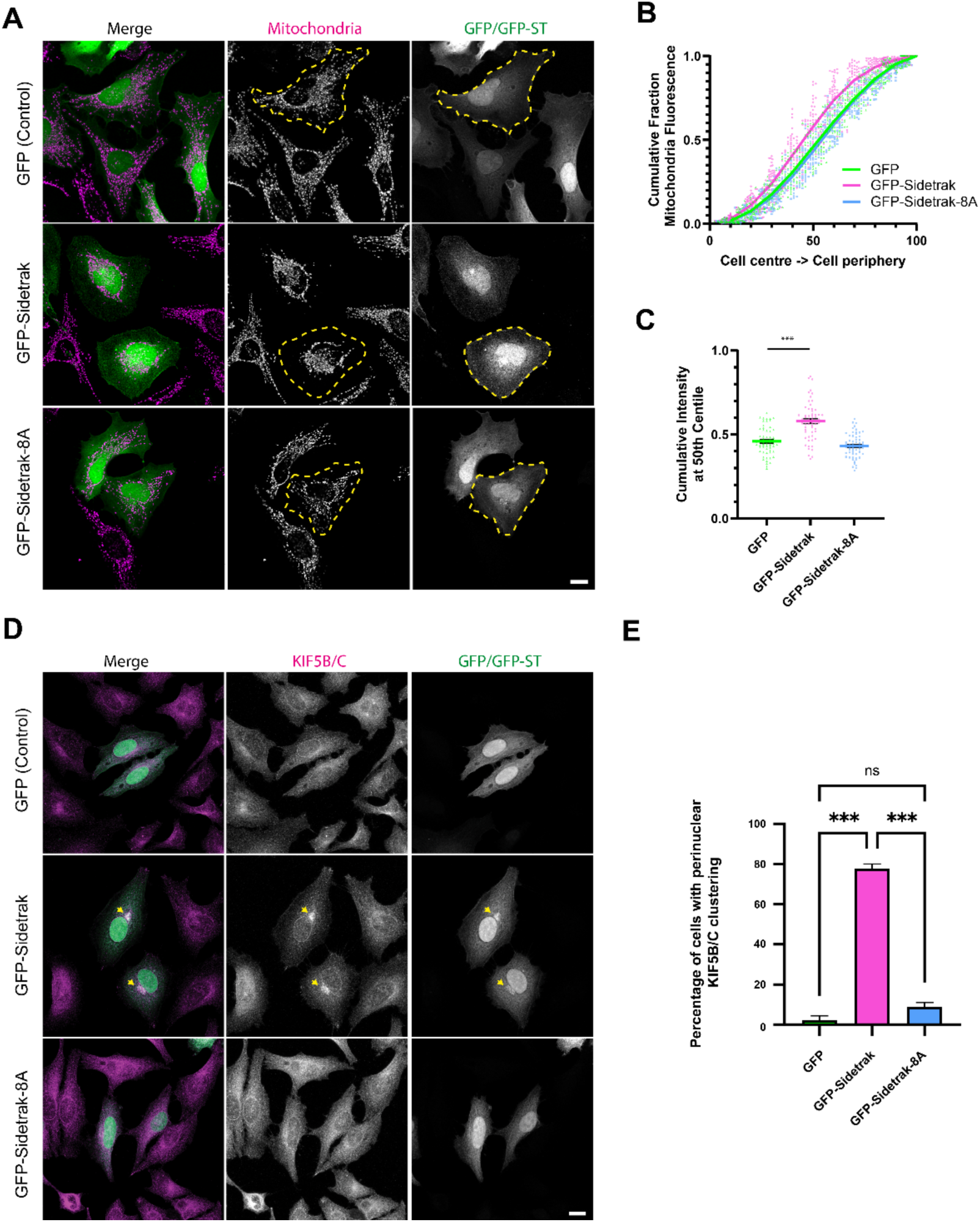
Sidetrak promotes perinuclear accumulation of mitochondria and kinesin-1. **(A)** Representative immunofluorescence images of cells expressing GFP, GFP-Sidetrak or the binding-defective GFP-Sidetrak-8A mutant, co-stained for mitochondria. **(B,C)** Quantification of mitochondrial distribution. Whole-cell regions of interest (ROIs) were defined and progressively scaled in 10% decrements towards the cell centre. The cumulative mitochondrial fluorescence intensity within each scaled ROI was measured and expressed as a fraction of the total mitochondrial fluorescence within the whole-cell ROI. Dashed yellow lines show cell boundary **(B)** Cumulative fraction of mitochondrial fluorescence as a function of radial extent, from the cell centre (10%) to the cell periphery (100%). **(C)** Fraction of total mitochondrial fluorescence contained within the central 50% of the cell ROI. Data represent 60 cells from three independent experiments and are shown as mean ± SEM. Statistical significance was determined by one-way ANOVA with Dunnett’s multiple-comparisons test comparing with GFP control. \*\*\**P* < 0.001. (D) Representative immunofluorescence images of cells expressing GFP, GFP-Sidetrak or GFP-Sidetrak-8A, co-stained for endogenous KIF5B/KIF5C. GFP-Sidetrak accumulates in perinuclear foci that are also positive for endogenous KIF5B/KIF5C (yellow arrows). **(E)** Quantification of perinuclear KIF5B/KIF5C accumulation. Data are from three independent experiments and are shown as mean ± SEM. Statistical significance was determined by one-way ANOVA with Tukey’s multiple-comparisons test comparing all pairs. \*\*\**P* < 0.001. Scale bars, 10 µm.

## Discussion

We have demonstrated that a solvent-exposed regulatory surface of a coiled-coil in kinesin-1 can be selectively targeted using *de novo* protein design. Starting from an AlphaFold3-derived model of the TPR-bound TDS, we designed Sidetrak, a nanomolar-affinity binder that selectively engages endogenous kinesin-1 complexes and inhibits their interaction with MAP7. These findings establish the TDS as a site for synthetic modulation and provide, to our knowledge, the first inhibitor of an interaction between a microtubule motor and a microtubule-associated protein. Thus, Sidetrak provides proof of concept that *de novo* protein design can manipulate endogenous motor function through a regulatory interface outside the conserved motor domain.

Coiled coils are widespread structural features in cytoskeletal motors and beyond, but present challenging targets for molecular recognition because their elongated geometry and repetitive surfaces provide relatively few distinctive features. However, coiled coils also frequently serve as regulatory interaction surfaces, providing interfaces through which multiple binding partners can modulate protein function. The kinesin-1 TDS exemplifies this principle, as it engages both the KLC TPR domains and MAP7, making it an established site of regulatory protein–protein interactions. Despite comprising just 36 amino acids, Sidetrak binds the exposed surface of a homodimeric coiled coil with high affinity and selectivity. This demonstrates that a compact designed binder can recognise a coiled-coil surface and suggests that coiled-coil-mediated regulatory interactions may provide targets for manipulation more broadly.

Our findings support a role for MAP7 function in maintaining mitochondrial distribution by directly engaging the motor complex. Sidetrak induces perinuclear mitochondrial redistribution, phenocopying genetic depletion of either MAP7 or kinesin-1 (*13*) indicating a net loss of kinesin-1 transport function following interference with MAP7 engagement in this context. The accompanying perinuclear accumulation of endogenous kinesin-1 is consistent with a shift in the balance between outward kinesin-1-dependent and inward dynein-dependent transport. Nonetheless, the molecular basis of these effects also requires consideration of the dual roles of the TDS. In our model, the TDS also contributes to autoinhibition through interaction with the KLC TPR domains (**Fig. 1A)**. Consistent with this, the kinesin-binding domain of MAP7 can promote some kinesin-1 activity independently of microtubule binding (*13, 20*) and MAP7 has also been reported to enhance kinesin-1 recruitment to cargoes by destabilising autoinhibition (*43*). By introducing a synthetic third mode of TDS occupancy, Sidetrak may impair microtubule recruitment while also altering kinesin-1 autoinhibition or cargo engagement. As such, it is best considered a modulator of this regulatory mechanism rather than an inhibitor of kinesin-1 activity *per se*. This distinction is consistent with our previous finding that small-molecule inhibition of cargo binding can promote kinesin-1 activity (*44, 45*). Importantly, Sidetrak binding may also alter the repertoire or stability of kinesin-1 interactions, potentially promoting associations that are otherwise transient or weakly populated. Thus, the Sidetrak-enriched proteome should not necessarily be considered a neutral representation of the endogenous kinesin-1 interactome, but rather a potentially modified interaction landscape arising from occupancy of this regulatory interface.

Sidetrak recognises all three kinesin-1 heavy-chain paralogues, consistent with complete conservation of its predicted binding site across KIF5A, KIF5B and KIF5C despite greater divergence in the surrounding CC1 sequences. This establishes Sidetrak as a pan-kinesin-1 binder and suggests a route to tuning the specificity of future designs: targeting conserved surfaces could enable family-wide modulation, whereas targeting divergent regions could confer paralogue selectivity. The choice of target site could also determine whether binding inhibits or activates motor function. In addition, such binders could be functionalised to promote cell-permeability and/or targeted protein degradation (*46, 47*). More broadly, coiled-coil regions outside the highly conserved motor domain offer potential targets for selectively manipulating other members of the kinesin superfamily. Similar strategies could extend to motors in the dynein and myosin families and their associated cargo adaptors, as well as other coiled-coil-containing proteins involved in membrane trafficking, such as tethering proteins.

Our findings also have implications for the local structure of CC1. AF3 predicts the TDS within a continuous canonical coiled coil, providing the *a priori* structural hypothesis used for Sidetrak design. The design template was generated from the AF3 model of the TPR-bound state by computational removal of the TPR domains, preserving the CC1 conformation predicted to accompany TPR engagement. Although the TPR domains were absent during binder generation, the predicted Sidetrak-binding mode resembles that of the first KLC TPR repeat. This convergence suggests recognition of the same CC1 surface and is consistent with the proposed TPR–TDS interaction in the autoinhibited state.

A recent cryo-EM study proposed a substantially different organisation of autoinhibited kinesin-1 (*20*). Although the KHC stalk is described as comprising coiled-coil segments, the deposited model (PDB: 9PMB) shows widespread departures from canonical packing, including separation of CC1. These departures accommodate the proposed overall architecture of the complex that is very different from AF3 models **(Fig. S1)** and contrast with structural and mutational evidence supporting canonical coiled-coil organisation of the neck-coil (CC0), the central elbow and the KHC–KLC interface (*5, 10, 48, 49*).

Sidetrak provides a direct test of CC1. It was designed against, and is predicted to engage, a composite epitope that alternates between the two helices of CC1, requiring them to be closely apposed **(Fig. S9).** Across the four-heptad TDS, mean inter-chain separation is 11.0 Å in the AF3-derived CC1/Sidetrak model, similar to canonical GCN4 (10.8 Å), but 17.1 Å in PDB: 9PMB **(Fig. S13).** Sidetrak nevertheless binds kinesin-1 with nanomolar affinity in a manner dependent on both the TDS and its designed interface, providing experimental evidence for this closely apposed CC1 geometry.

Formally, Sidetrak could selectively recognise a canonically packed open state of kinesin-1. However, the proposed autoinhibited architecture requires disruption of coiled-coil packing across much of the stalk, including the high-affinity KHC–KLC interface, whose energetic cost would need to be compensated by interactions formed specifically in the closed state. Given that kinesin-1 autoinhibition is a readily shifted equilibrium (*5, 10, 50*) we favour a model where canonical coiled-coil packing provides the underlying structural framework in both states, with localised regulatory interactions sufficient to shift the equilibrium between them.

In summary, Sidetrak demonstrates that a motor–MAP interaction can be disrupted by targeting a regulatory coiled-coil surface. Its high-affinity binding also illustrates the potential of *de novo* protein design to generate compact ligands that recognise relatively feature-poor coiled-coil surfaces. Designed binders that selectively modulate cytoskeletal motors may provide new tools to dissect their structures, functions and mechanisms and, ultimately, to address disease-associated transport defects.

## Acknowledgements

This work was funded by UKRI-BBSRC grant BB/Z517276/1 and MPD acknowledges additional support from a Lister Institute of Preventative Medicine Fellowship. JAC was supported by a Wellcome Trust Early Career Award (328547/Z/25/Z) and Academy of Medical Sciences Springboard Award (SBF0011\1170). TB and SAB were supported by the Wellcome Trust PhD Programme in Dynamic Molecular Cell Biology at the University of Bristol (218510/Z/19/Z). DNW is supported by the Novo Nordisk Foundation through the NNF Center for Protein Design at the University of Copenhagen (NNF25SA0105927). We are grateful to Dr Kate Heeson and Dr Phil Lewis at the University of Bristol Proteomics Facility for support with mass-spectrometry and analysis. We thank Roberto Steiner (King’s College London/University of Padova) for helpful discussions.

## Author contributions

Investigation: J.A.C., S.A.B., M.P.D; Resources: J.A.C., T.B., M.P.D.; Conceptualization: J.A.C., D.N.W., M.P.D.; Methodology: M.P.D.; Supervision: M.P.D.; Formal analysis: M.P.D.; Writing – original draft: M.P.D.; Writing – review & editing: J.A.C., S.A.B., T.B., D.N.W., M.P.D.

## Data Availability

The complete mass spectrometry proteomics dataset, including raw data files, will be deposited in the ProteomeXchange Consortium via the PRIDE partner repository and made publicly available upon publication. The dataset accession number will be provided once assigned.

## Materials and Methods

### AlphaFold3 Modelling

All models shown were generated using the AlphaFold3 webserver (alphafoldserver.com)(*21*). PAE plots are as presented by the software and manually annotated. Where indicated, the 5 output models were reranked using FoldScript (*51*), and pLDDT is shown using the standard colour scheme. Protein sequences (accession numbers and amino acid ranges) are provided in the figure legends.

### Boltzgen binder design

BoltzGen was downloaded and installed from its GitHub repository (https://github.com/HannesStark/boltzgen) and run on a Linux workstation equipped with an AMD Ryzen Threadripper PRO CPU and two NVIDIA RTX 5000 Ada Generation GPUs. The design workflow, sequences/models used and target hotspots are described in **Fig. S3.**

### Plasmid DNA

Binders were supplied by GenScript as mammalian cell codon-optimised synthetic DNA constructs in pcDNA3.1(+)-N-eGFP. Binder sequences were cloned downstream of eGFP using KpnI and EcoRI restriction sites, with protein sequences as shown in Fig. S3. A gly–ser linker was positioned immediately N-terminal to each binder. Together with the amino acids encoded by the KpnI cloning site, this generated a four-residue gly-thr-gly-ser linker between eGFP and the binder (GFP-*GTGS*-Binder). Amino acid sequence of Sidetrak is: SERLRELRARLERARARGDETLVRALELVIRYLEES. Sidetrak-8A was also obtained as synthetic DNA in an identical backbone. Mammalian cell kinesin-1 expression constructs have been described previously (*12, 52*). His-K560, comprising the first 560 amino acids of rat KIF5C, was subcloned into the bacterial expression vector pMW-His-3C using NotI and EcoRI restriction sites. The shorter K401 construct was generated by PCR and subcloning. mCherry-MAP7 was generated by PCR amplification of MAP7 from Addgene plasmid 46076 (pEGFP-MAP7) (*14*) and subcloning into pmCherry-C1 using HindIII and BamHI restriction sites. All plasmids were verified by DNA sequencing.

### Cell culture and transfection

All cells were maintained in Dulbecco’s modified Eagle’s medium (DMEM) with GlutaMAX-I (Invitrogen), supplemented with 10% heat-inactivated fetal bovine serum (HI-FBS), 100 U/ml penicillin and 100 µg/ml streptomycin, at 37°C in a 5% CO₂ atmosphere. Transfections were carried out using Effectene transfection reagent (Qiagen) according to the manufacturer’s instructions.

### Immunofluorescence and cell imaging

Cells were grown on fibronectin-coated coverslips in 6-well plates and transfected with the indicated constructs. At 16 h post-transfection, cells were fixed with 4% paraformaldehyde for 15 min at room temperature and permeabilized with 0.1% Triton X-100 in PBS for 2 min. Cells were blocked in PBS supplemented with 3% BSA and incubated with primary and secondary antibodies diluted in blocking buffer for 30 min. Anti-HA antibody (HA-7, H3663; Sigma-Aldrich/Merck), anti-mitochondria antibody (ab92824; Abcam) and anti-KIF5B/5C antibody (ab167429; Abcam) were each used at a 1:1,000 dilution and detected using Alexa Fluor 568- or Alexa Fluor 647-conjugated anti-mouse secondary antibodies. Imaging was performed using a Leica SP8 confocal laser-scanning microscope (Leica Microsystems, Germany) equipped with a 63×/1.4 NA oil-immersion objective. Images, shown as maximum-intensity projections unless otherwise indicated, were prepared for publication using ImageJ/Fiji and Adobe Photoshop/Illustrator.

### Protein production and purification

An expression plasmid encoding His-tagged K560 (rat KIF5C, residues 2-560) was transformed into BL21(DE3) cells. Cells were cultured in LB Miller medium with shaking at 37°C and 180 rpm. When the optical density at 600 nm (OD₆₀₀) reached approximately 0.6, the temperature was reduced to 18°C and protein expression was induced by addition of 0.2 mM IPTG. Cultures were incubated overnight with shaking at 18°C. Cells were harvested by centrifugation at 5,000 × *g* for 15 min at 4°C and resuspended in Buffer A (40 mM HEPES, pH 7.4, 500 mM NaCl, 40 mM imidazole, 5% glycerol and 5 mM β-mercaptoethanol) supplemented with protease inhibitors. Cells were lysed by sonication (70% amplitude, 5 s on/15 s off for a total of 7.5 min) on ice. Lysates were clarified by centrifugation at 40,000 × *g* for 45 min at 4°C, and the resulting supernatant was filtered through a 0.45 µm membrane. The clarified lysate was loaded onto a 1 ml HisTrap column equilibrated with Buffer A. The column was washed with at least 50 column volumes of Buffer A containing 30 mM imidazole, and bound protein was eluted using a gradient to 500 mM imidazole. Eluted fractions were analysed by SDS–PAGE, and fractions containing His-K560 were pooled and concentrated using a 10 kDa molecular-weight cut-off centrifugal concentrator to approximately 500 µl before size-exclusion chromatography (SEC). SEC was performed using a Superose 6 10/300 column equilibrated in 20 mM HEPES, pH 7.4, 150 mM NaCl, 1 mM MgCl₂, 0.1 mM ADP and 1mM β-mercaptoethanol. Eluted fractions were analysed by SDS–PAGE, and fractions containing purified His-K560 were pooled, aliquoted, flash-frozen and stored at −80°C until use. Protein concentration was determined using a spectrophotometer and confirmed by running on a gel using BSA standards.

### Fluorescence polarization binding assays

TAMRA-conjugated Sidetrak peptide was diluted to 100 nM and incubated with increasing concentrations of purified K560/K401 in assay buffer (25 mM HEPES pH 7.4, 150 mM NaCl, 5 mM β-mercaptoethanol). Measurements were performed at room temperature on a POLARStar Omega microplate reader (BMG Labtech). K560/K401 and Sidetrak concentrations are expressed per monomer. AF3 modelling predicts two equivalent, non-contacting Sidetrak sites per CC1 dimer, related by the C2 axis, so the molar concentration of binding sites equals the monomer concentration. Polarisation values were corrected by subtraction of peptide-alone controls, normalised to the fitted Bmax, and fitted to a quadratic one-site tight-binding equation accounting for ligand depletion (*4, 53*). Data are from triplicate samples and are representative of three independent experiments.

### GFP-TRAP immunoprecipitation and western blotting

HEK293 cells grown in 10 cm dishes were plated at 50% confluency and transfected with the indicated plasmids. At 16 h post-transfection, the medium was removed and cells were washed with wash buffer (25 mM HEPES, pH 7.4, 150 mM NaCl). Cells were lysed in 800 µl lysis buffer (wash buffer supplemented with 0.1% NP-40, 0.1% Triton X-100 and protease inhibitor cocktail), and lysates were clarified by centrifugation at 13,000 × *g* for 10 min. A 40 µl sample of each lysate was retained as the input and mixed with 4× SDS loading buffer before incubation at 95°C for 5 min. GFP-TRAP beads (25 µl per sample) were washed three times in wash buffer before addition to the clarified cell extracts, and immunoprecipitation was performed for 1.5 h at 4°C with mixing. In experiments where Sidetrak peptide was used as a competitor, the peptide was added to the clarified lysates and incubated for 15 min before addition of the GFP-TRAP beads. Control and peptide-treated samples contained 0.2% DMSO.

In all cases, beads were washed three times with wash buffer before resuspension in 1× SDS loading buffer and incubation at 95°C for 5 min. Aliquots (15 µl) of input and immunoprecipitated samples were resolved by SDS-PAGE prior to transfer and western blotting. Membranes were blocked in PBS containing 5% milk and 0.1% Tween-20 prior to incubation with the indicated primary antibodies. Anti-HA antibody (HA-7, H3663; Sigma-Aldrich/Merck), anti-Myc antibody (9E10; Thermo Fisher Scientific) and anti-GFP antibody (clones 7.1 and 13.1, 11814460001; Roche) were each used at a 1:1,000 dilution. Goat anti-mouse Alexa Fluor 680 secondary antibody (A21057; Invitrogen) was used at a 1:10,000 dilution. Images were acquired and quantified using a LI-COR Odyssey CLx imaging system and Image Studio software.

### Quantification of mitochondrial distribution

Mitochondrial distribution was quantified in Fiji. Whole-cell regions of interest (ROIs) were manually defined using the cell boundary visualised in the GFP channel and progressively scaled towards the centre of the ROI in 10% increments, generating nested ROIs corresponding to 10–100% of the linear dimensions of the original whole-cell ROI. Integrated mitochondrial fluorescence intensity was measured from background-subtracted images within each ROI and expressed as a fraction of the total mitochondrial fluorescence intensity within the original whole-cell ROI. Cumulative radial distribution profiles were generated by plotting the fraction of total mitochondrial fluorescence against ROI scale. The fraction of mitochondrial fluorescence contained within the central 50% ROI was used as a single-cell measure of mitochondrial redistribution. Measurements were performed on 60 cells from three independent experiments. Statistical analyses were performed in GraphPad Prism using one-way ANOVA with Dunnett’s multiple-comparisons test.

### Peptide synthesis, purification and characterisation

Sidetrak peptides were synthesised by automated Fmoc solid-phase synthesis (CEM Liberty Blue) on a 0.1mM scale using, with N-terminal acetylation or TAMRA labelling, as previously described (*46*). Activation was achieved with DIC/Oxyma. Fmoc deprotection was performed with 20% v/v morpholine/dimethyl formamide (DMF). Synthesis was from C to N terminus as the C-terminal amide on Rink amide resin. Labelling was achieved by addition of TAMRA (0.1 mM, 2 equiv.), HATU (0.095 mM, 1.9 equiv.) and DIPEA (0.225 mM, 4.5 equiv.) in DMF (3 ml) to DMF washed peptide resin (0.05 mM) with agitation for 3 hours. Resin was washed with 20% piperidine in DMF (5 ml) for 2 × 30 minutes to remove any excess dye. All manipulations were carried out under foil to exclude light. Alternatively, N-terminally acetylated peptide was produced by addition of acetic anhydride (0.25 ml) and pyridine (0.3 ml) in DMF (5 ml) for 30 min at room temperature. Peptides were cleaved from the solid support by addition of TFA (9.5 ml), TIPS (0.25 ml) and water (0.25 ml) for 3 hours with shaking at room temperature. The cleavage solution was reduced to approximately 1 ml under a flow of nitrogen. Crude peptide was precipitated upon addition of ice-cold diethyl ether (40 ml) and recovered via centrifugation. The resulting precipitant was dissolved in 1:1 acetonitrile and water (∼15 ml) and lyophilized to yield crude peptide as a solid.

Peptides were purified by reverse-phase high-performance liquid chromatography (HPLC) on a Phenomenex Luna C18 stationary phase column (150 × 10 mm, 5-µm particle size, 100-Å pore size) using a preparative JASCO HPLC system. A linear gradient of 20–80% acetonitrile and water (with 0.1% trifluoroacetic acid) was applied over 30 minutes. The identities of the peptides were confirmed using matrix-assisted laser desorption/ionisation–time-of-flight (MALDI–TOF) mass spectrometry using a Bruker ultrafleXtreme II instrument. Peptides were spotted on a ground-steel target plate using dihydroxybenzoic acid or α-cyano-4-hydroxycinnamic acid as the matrix. Peptide concentrations were determined by UV–visible absorbance using sequence-derived extinction coefficients at 280 nm or, for TAMRA-labelled peptides, an extinction coefficient of 85,000 M⁻¹ cm⁻¹ at 555 nm.

### Structural comparison of CC1 models

The geometry of the TPR docking site (TDS) was compared between the AF3-derived CC1/Sidetrak design model and the deposited cryo-EM model of kinesin-1 (PDB: 9PMB). Corresponding four-heptad regions comprising the TDS of CC1 were analysed. For each model, pairwise inter-chain distances were calculated between Cβ atoms for all residues within the selected region, generating a 28 × 28 distance matrix. Differences between corresponding distance matrices were quantified as the root-mean-square (RMS) deviation, mean absolute deviation and maximum absolute deviation in inter-chain distance. Distances between equivalent residue positions across the two CC1 chains were additionally analysed separately.

### Tandem mass tag proteomic analysis

Immuno-isolated samples (from GFP-TRAP pull downs) were reduced with 10 mM TCEP at 55°C for 1 h, alkylated with 18.75 mM iodoacetamide at room temperature for 30 min, and digested on the beads with 2.5 µg trypsin at 37°C overnight. The resulting peptides were labelled with TMT ten-plex reagents according to the manufacturer’s protocol (Thermo Fisher Scientific, Loughborough, UK). Labelled samples were pooled and desalted using a SepPak cartridge according to the manufacturer’s instructions (Waters, Milford, Massachusetts, USA). The eluate was evaporated to dryness and resuspended in 1% formic acid before analysis by nano-LC–MS/MS using an Ultimate 3000 nano-LC system coupled to an Orbitrap Fusion Tribrid mass spectrometer (Thermo Scientific).

Peptides in 1% (vol/vol) formic acid were injected onto an Acclaim PepMap C18 nano-trap column (Thermo Scientific). After washing with 0.5% (vol/vol) acetonitrile and 0.1% (vol/vol) formic acid, peptides were resolved on a 250 mm × 75 µm Acclaim PepMap C18 reversed-phase analytical column (Thermo Scientific) over a 150 min organic gradient at a flow rate of 300 nl min⁻¹. The gradient comprised seven segments: 1–6% solvent B over 1 min, 6–15% B over 58 min, 15–32% B over 58 min, 32–40% B over 5 min, 40–90% B over 1 min, 90% B for 6 min, and a reduction to 1% B over 1 min. Solvent A was 0.1% formic acid, and solvent B was aqueous 80% acetonitrile in 0.1% formic acid. Peptides were ionised by nano-electrospray ionisation at 2.0 kV using a stainless-steel emitter with an internal diameter of 30 µm (Thermo Scientific) and a capillary temperature of 275°C.

Spectra were acquired using an Orbitrap Fusion Tribrid mass spectrometer controlled by Xcalibur 2.1 software (Thermo Scientific), operated in data-dependent acquisition mode using an SPS-MS3 workflow. FTMS1 spectra were collected at a resolution of 120,000, with an automatic gain control (AGC) target of 200,000 and a maximum injection time of 50 ms. Precursors were filtered using an intensity threshold of 5,000 and charge states of 2–7, with monoisotopic peak determination set to peptide. Previously interrogated precursors were excluded using a dynamic window of 60 s ±10 ppm. MS2 precursors were isolated using a quadrupole isolation window of 1.2 m/z. ITMS2 spectra were collected with an AGC target of 10,000, a maximum injection time of 70 ms and a CID collision energy of 35%. For FTMS3 analysis, the Orbitrap was operated at a resolution of 50,000, with an AGC target of 50,000 and a maximum injection time of 105 ms. Precursors were fragmented by higher-energy collisional dissociation (HCD) at a normalised collision energy of 60% to ensure maximal TMT reporter-ion yield. Synchronous precursor selection (SPS) was enabled to include up to ten MS2 fragment ions in the FTMS3 scan.

### Quantitative proteomic data analysis

Mass spectrometry data were searched against the UniProt *Homo sapiens* database downloaded on 18 January 2026, supplemented with the sequences of the experimental constructs and a common contaminants database. Protein groups were assigned using Proteome Discoverer version 2.4. Protein accession annotations were updated against UniProt on 9 October 2026, and representative master proteins were selected from the candidates within each group according to their current review and annotation status, without altering the underlying identification or quantification evidence. Three independent immunoprecipitates were analysed per condition: GFP-Sidetrak, GFP-Sidetrak-8A and GFP alone. All samples were prepared together, and the experimental design was unpaired. Downstream analyses were performed in R version 4.5.1. Protein abundances were log₂-transformed and normalised to median protein abundance. Median normalisation was selected following exploratory assessment of abundance distributions, within-condition variability and principal component analysis. A single linear model incorporating all three conditions was fitted using LIMMA, without pairing or blocking terms. Contrasts were specified for GFP-Sidetrak versus GFP, GFP-Sidetrak versus GFP-Sidetrak-8A, and GFP-Sidetrak-8A versus GFP. Statistical significance was assessed using empirical Bayes-moderated t-tests, and P values were adjusted for multiple testing using the Benjamini–Hochberg false discovery rate (FDR) method. Proteins were classified as enriched or depleted at FDR <0.05 and an absolute log₂ fold change >1. Results presented were restricted to protein groups supported by at least three unique peptides. Volcano plots display log₂ fold changes against −log₁₀ unadjusted P values, with adjusted significance indicated separately. MA plots display log₂ fold changes against mean log₂ abundance. For heatmap visualisation, log₂ abundances of selected proteins were standardised to Z-scores within each protein across samples.

## Supplementary Figures

**Figure S1.**
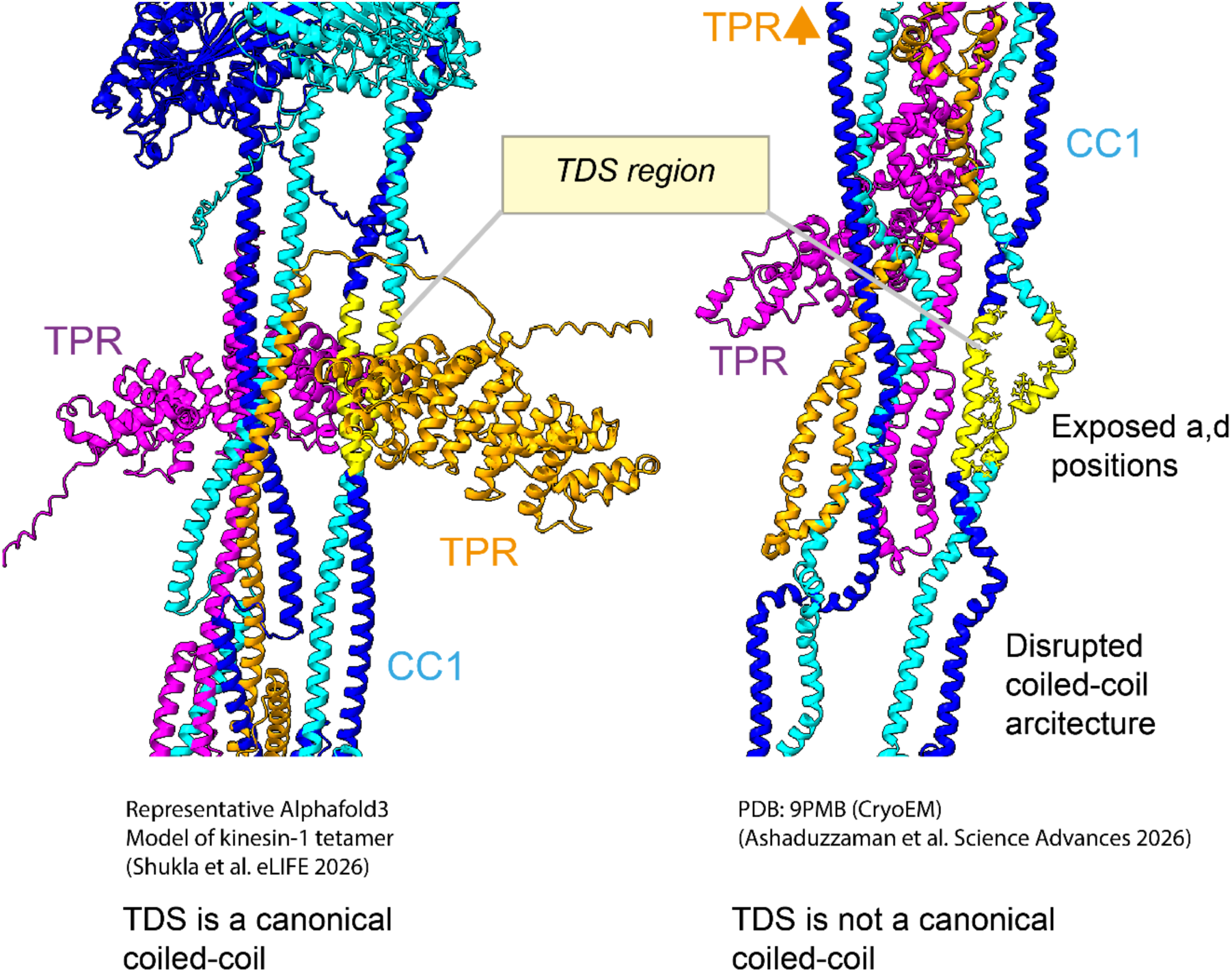
Comparison of AlphaFold3 model and Cryo-EM derived model for the TDS and surrounding coiled coils. Left, representative AlphaFold3 (AF3) model, as described in (*12*), in which the kinesin light chain TPR domains dock at the TDS (yellow). The TDS and surrounding CC1 sequences form a canonical heptad-based parallel coiled-coil dimer. Right, atomic model derived from the cryo-EM density map (PDB:9PMB)(*20*). In this model, CC1 does not adopt a canonical coiled-coil architecture: the helices comprising the TDS and elsewhere are widely separated, and the TPR domains occupy alternative positions. One TPR domain is shown, the other engages the motor domains.

**Figure S2.**
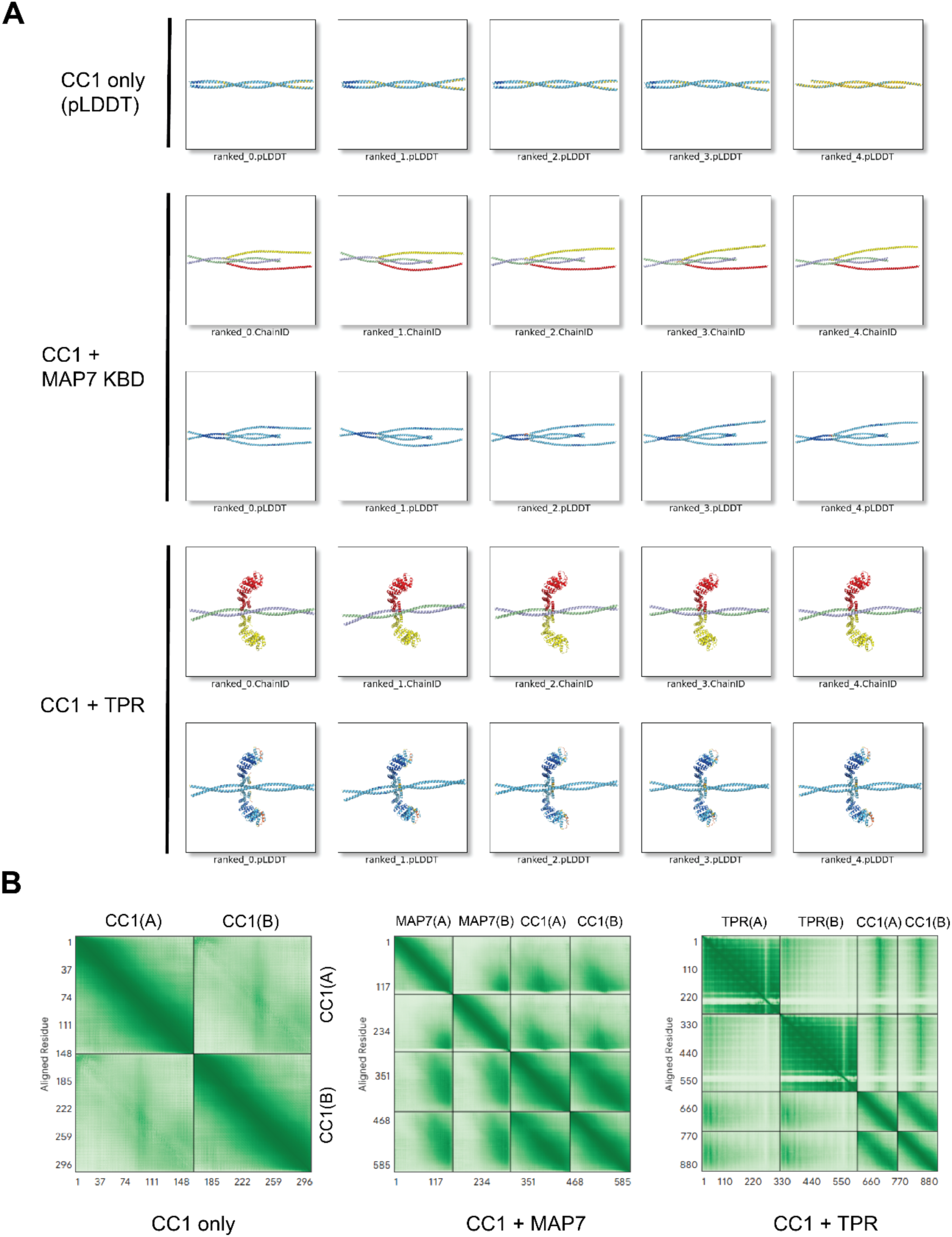
AF3 models of CC1 alone and in complex with MAP7 or TPR domains. AF3 models of KIF5C CC1 alone (NP_001101200.1; a.a. 410–558), in complex with the KLC TPR domains (NP_032476.2; a.a. 205–495), or with the kinesin-binding domain of MAP7 (NP_001185564.1; a.a. 467–610). The five AF3 models generated for each complex were reranked using FoldScript and are coloured by pLDDT or protein chain. The top-ranked models correspond to those shown in Fig. 1B. **(B)** PAE plots for the top-ranked models, annotated by chain.

**Figure S3.**
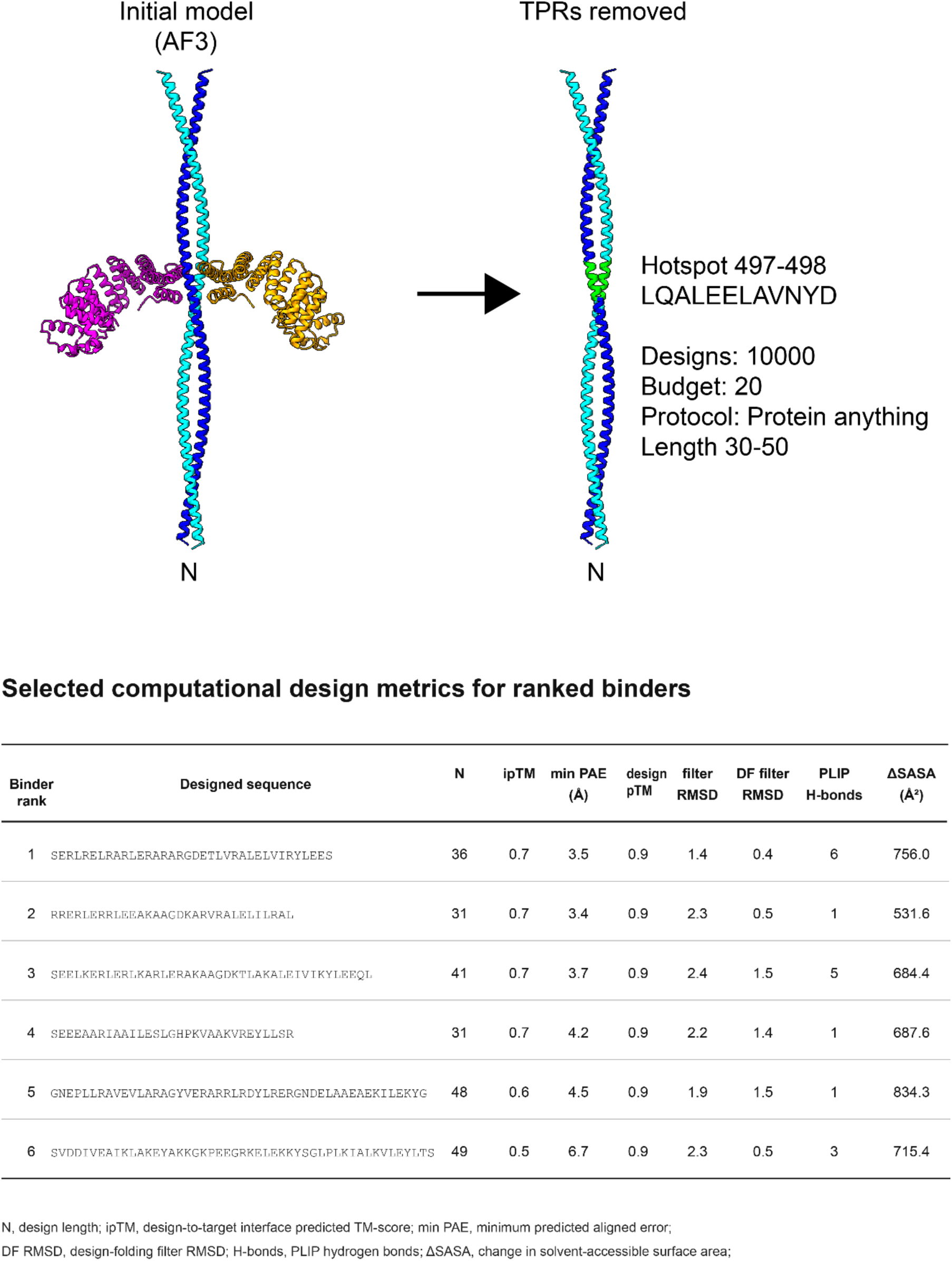
BoltzGen design workflow and output. Summary of BoltzGen design workflow and selected metrics for the six candidate binders from 10,000 designs. Initial model was the top ranked model with sequences as described in Fig. S2. All six candidates passed the default BoltzGen filters and were taken forward for further computational and experimental characterisation. Reported metrics were calculated for the 2:1 CC1–binder complexes (2× CC1 chains and 1× binder) used in the design pipeline.

**Figure S4.**
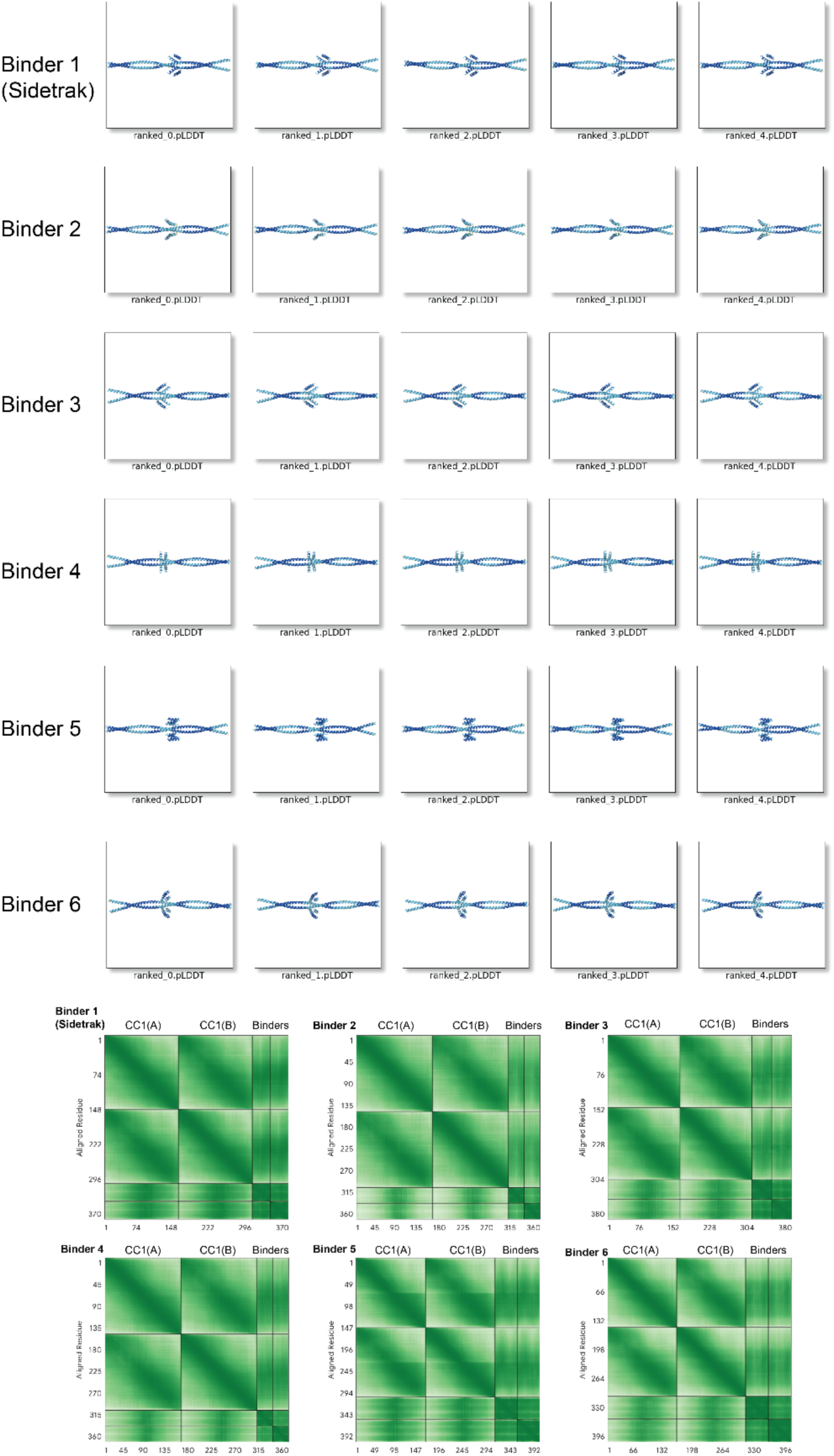
AF3 models of 6 candidate binders as 2:2 heterotetramers with CC1. AF3 models of KIF5C CC1 (NP_001101200.1; a.a. 410–558), in complex with 6 candidate binders (sequences as shown in Fig. S3). Modelled as 2:2 heterotetramers and reranked using FoldScript. Models are coloured by pLDDT. **(B)** PAE plots for the top-ranked models, annotated by chain.

**Figure S5.**
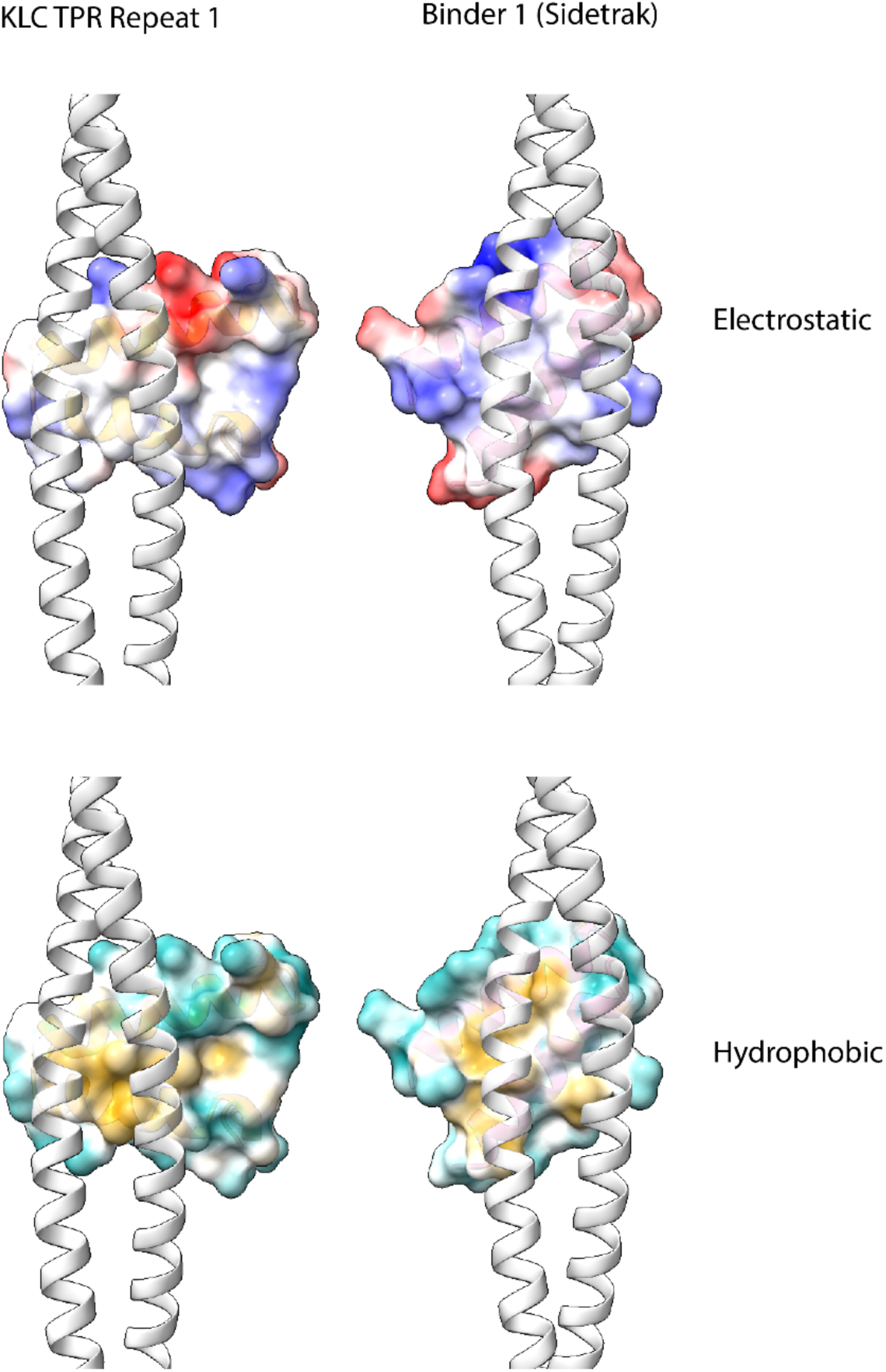
Comparison of surface properties of KLC TPR1 and Sidetrak at the predicted interface with CC1. AF3 models showing electrostatic and hydrophobic surface representations of KLC TPR1 and Binder 1 (Sidetrak) generated in UCSF Chimera X at their predicted interfaces with the CC1 TDS. KLC TPR (Repeat 1) is shown isolated from the full KLC TPR–CC1 model for clarity.

**Figure S6.**
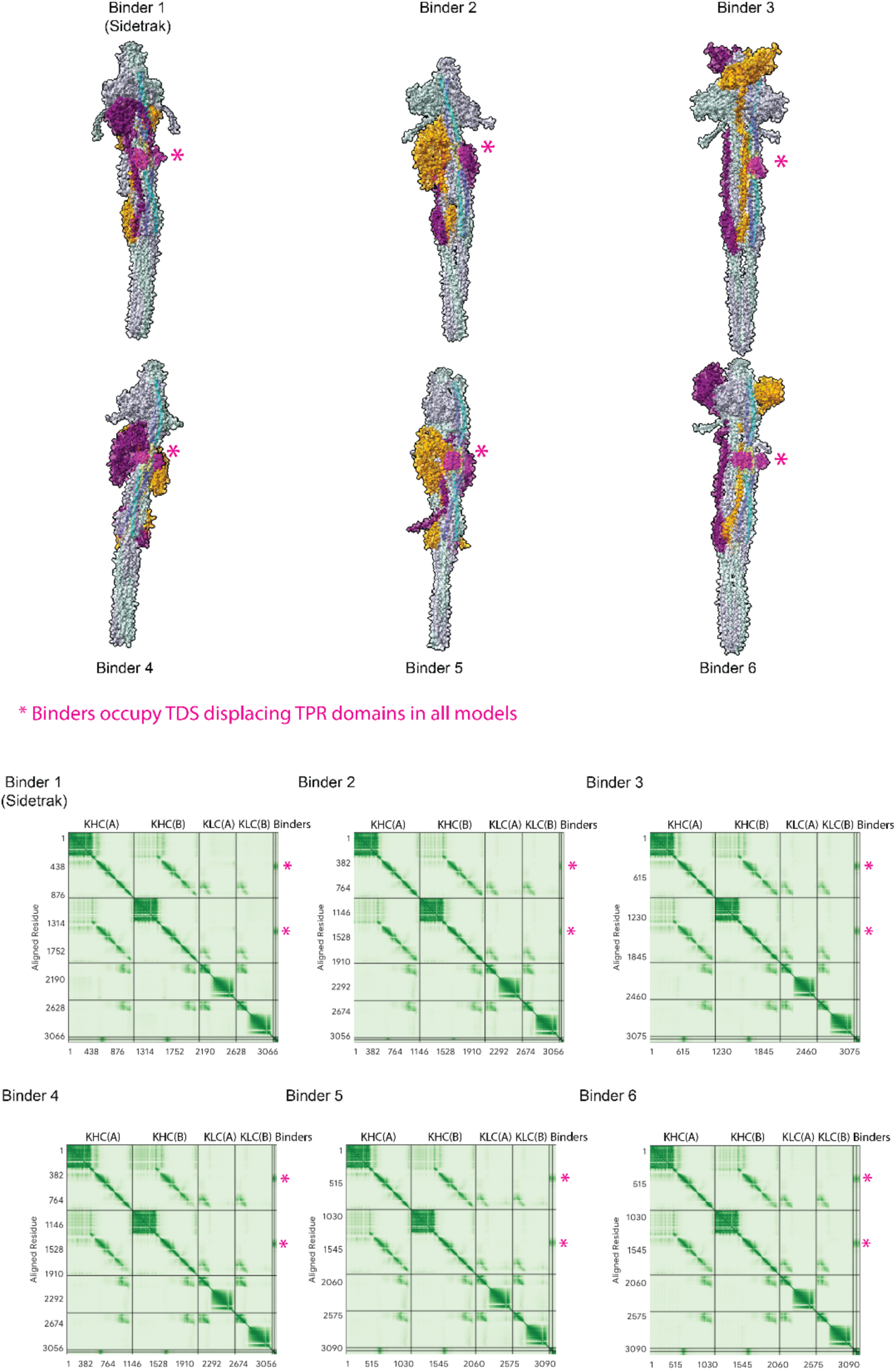
AF3 models of six candidate binders in complex with the complete kinesin-1 heterotetramer. (A) AF3 models of each of the six candidate binders in complex with the complete kinesin-1 heterotetramer, comprising the KHC motor domains and coiled-coil scaffold together with the coiled coil and TPR domains of KLC. Binders were modelled at the CC1 TDS and are shown in magenta. (B) PAE plots for the corresponding top-ranked models, annotated by chain. * indicates position of binding and low PAE of binder vs KHC.

**Figure S7.**
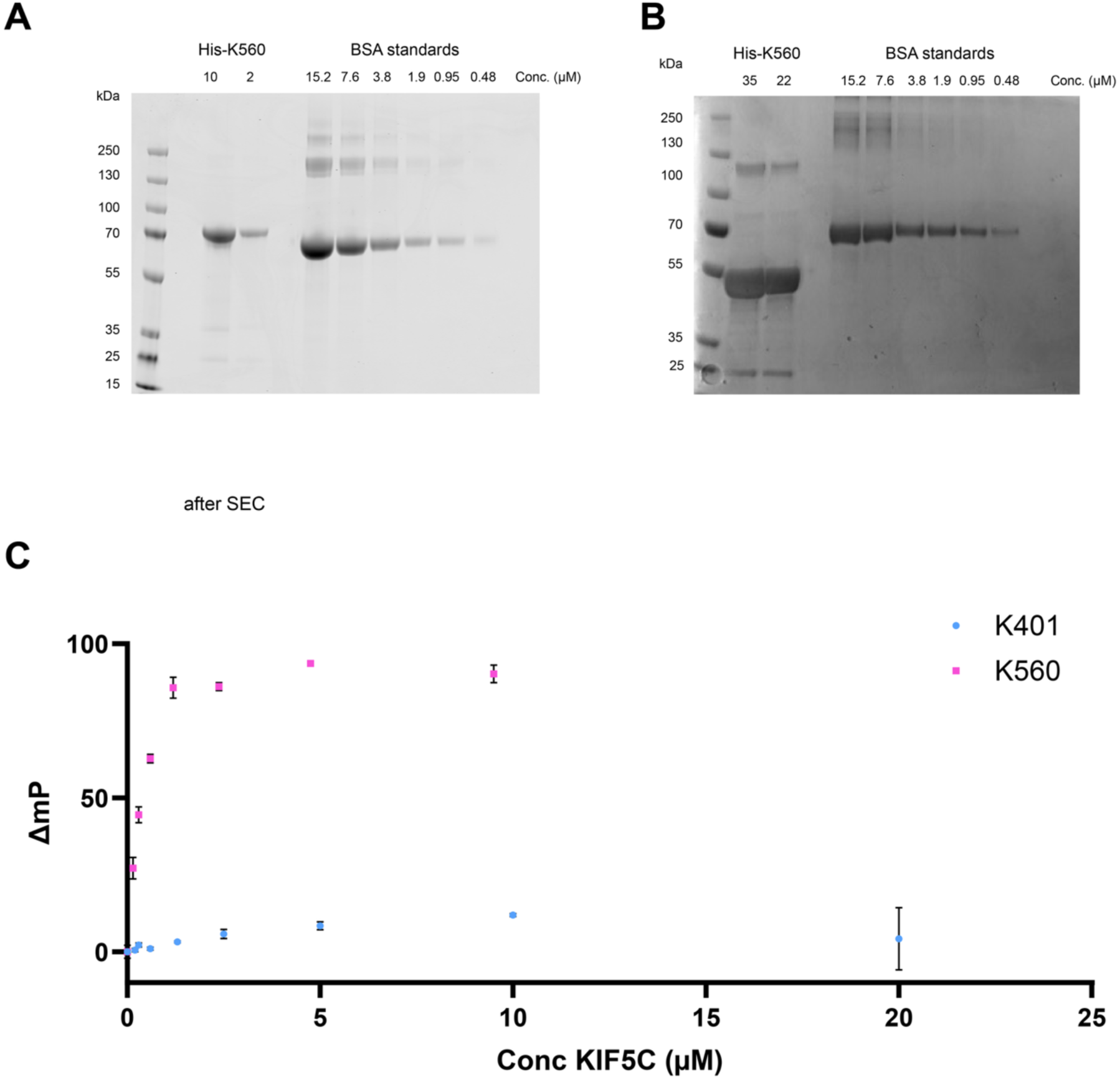
Protein samples and data for fluorescence polarisation experiments. (A, B) Coomassie-stained SDS–PAGE gels showing samples of K560 and K401 (His-KIF5C 2–560 and His-KIF5C 2–401, respectively) following size-exclusion chromatography. (C) Fluorescence polarisation measurements of TAMRA-Sidetrak in the presence of the indicated concentrations of K560 or K401. Data are presented without normalisation or curve fitting. The corresponding normalised and fitted data for Sidetrak/K560 are presented in Fig. 2.

**Figure S8.**
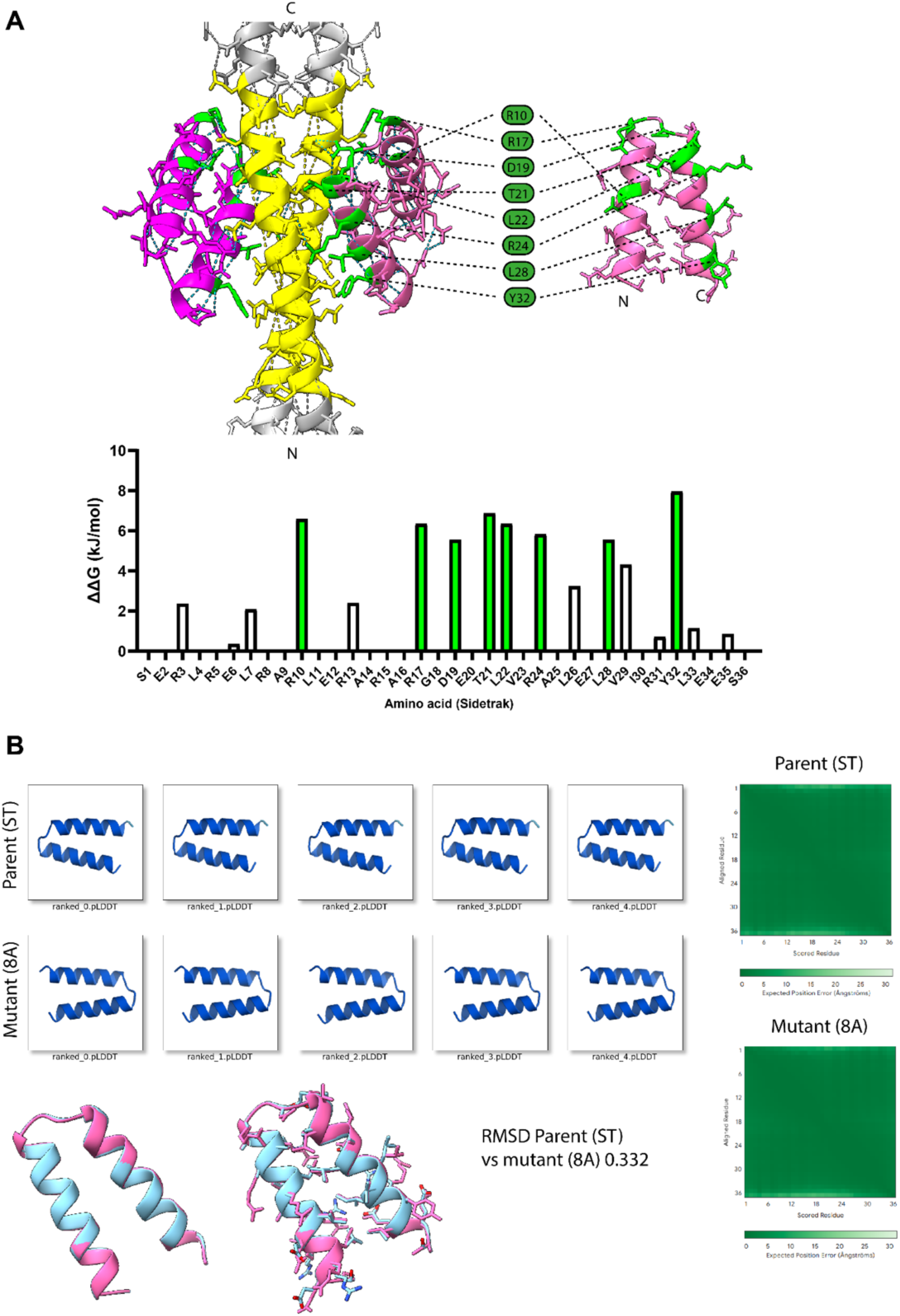
Design of mutations in Sidetrak to disrupt TDS binding. (A) AF3 model of Sidetrak bound to the KIF5C CC1 TDS, highlighting Sidetrak residues predicted to contribute to the binding interface using BUDE computational alanine scanning of Sidetrak. Graph showing the predicted energetic contribution (ΔΔG) of individual residues to the interaction with CC1. Interface residues with predicted ΔΔG ≥ 5 kJ/mol are shown in green were selected for simultaneous substitution with alanine to generate the Sidetrak-8A mutant. (B) Comparison of AF3 models of wild-type Sidetrak and Sidetrak-8A. The introduced alanine substitutions are confined to the predicted CC1-binding surface and do not disrupt the hydrophobic seam of the α-helical hairpin. Both proteins are predicted to adopt essentially identical overall structures, consistent with selective disruption of the designed binding interface.

**Figure S9.**
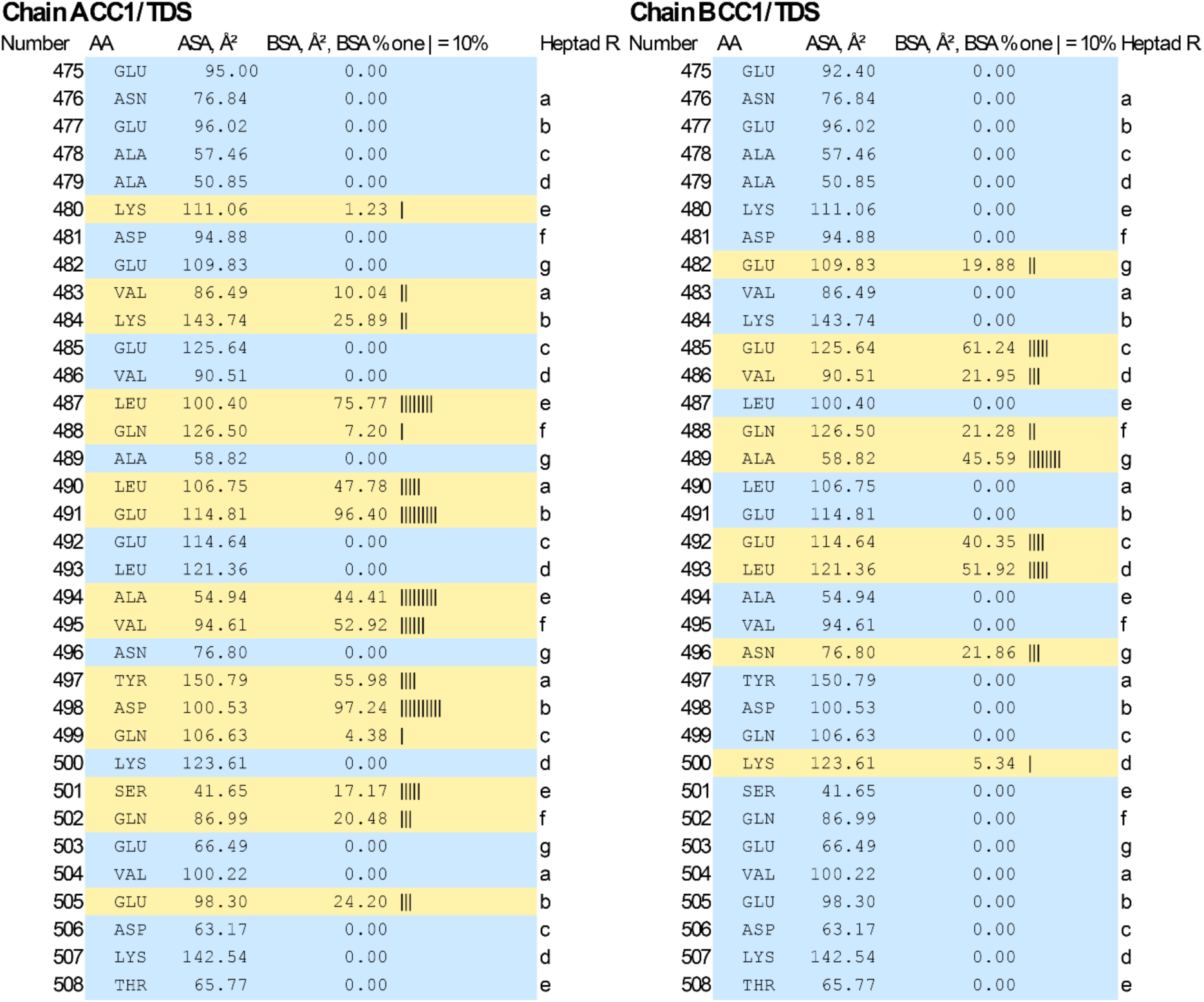
PISA analysis of the Sidetrak-binding surface on the KIF5C CC1/TDS coiled coil. Residue-level solvent-accessible surface area (ASA) and buried surface area (BSA) were calculated for chains A and B of the KIF5C CC1/TDS coiled coil following interaction with a single Sidetrak molecule. BSA is shown in Å² together with a graphical representation of the proportion of each residue’s ASA buried at the interface, where each vertical bar represents 10% burial. Heptad register assignments (a–g) are shown for each residue. The analysis reveals a composite Sidetrak-binding surface formed by both helices of the KIF5C coiled coil, with interacting residues from chains A and B distributed across the same region of CC1/TDS.

**Figure S10.**
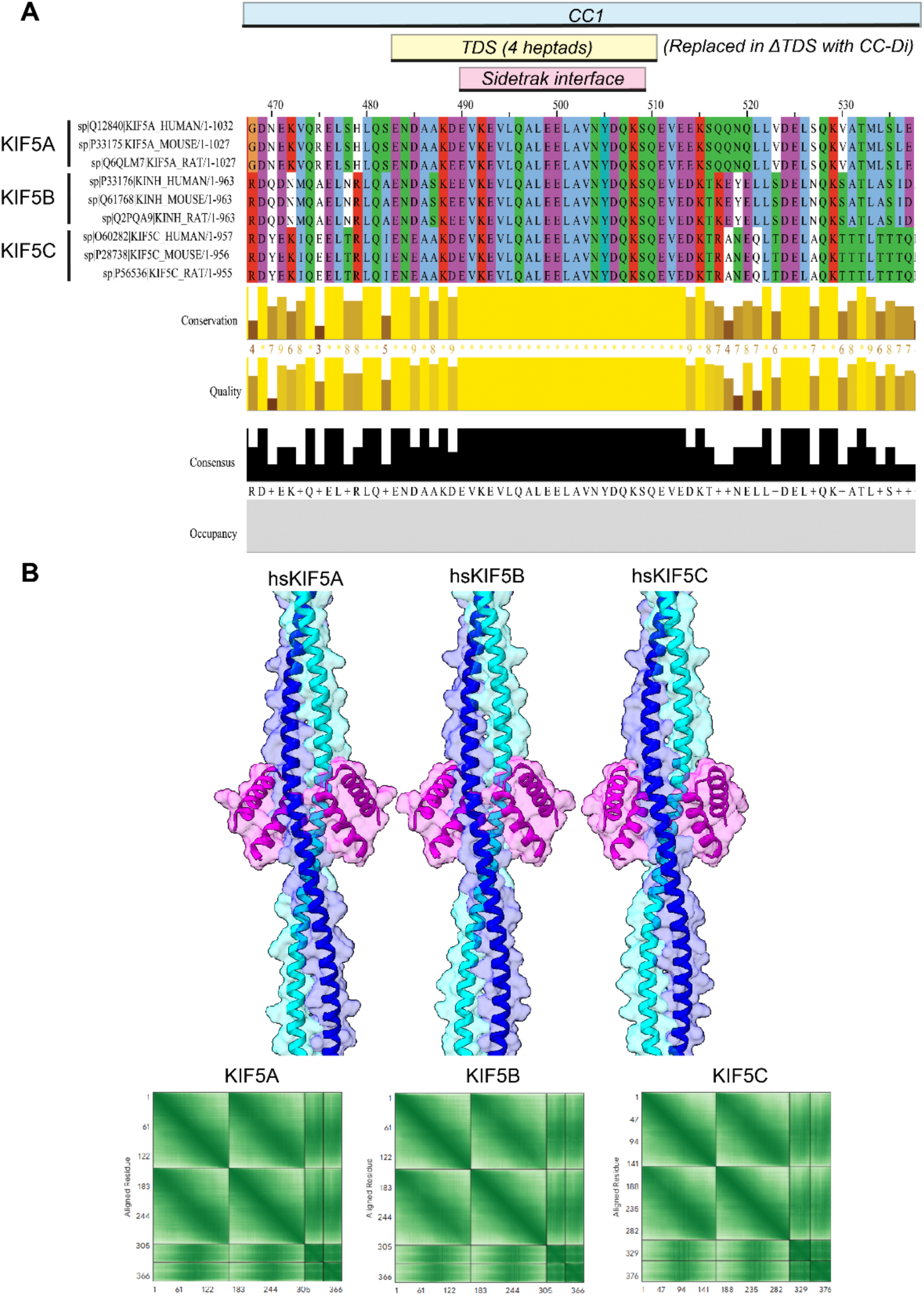
Sequence conservation and structural modelling of Sidetrak-TDS interaction across KIF5 paralogues. (A) Multiple sequence alignment (MAFFT, visualised using Jalview (*55*)) of human, mouse and rat orthologues of KIF5A, KIF5B and KIF5C, centred on the TDS. Sequences and accession numbers are indicated in the figure. (B) AF3 models of Sidetrak in complex with the CC1 TDS of human KIF5A, KIF5B and KIF5C. PAE plots are shown below.

**Figure S11.**
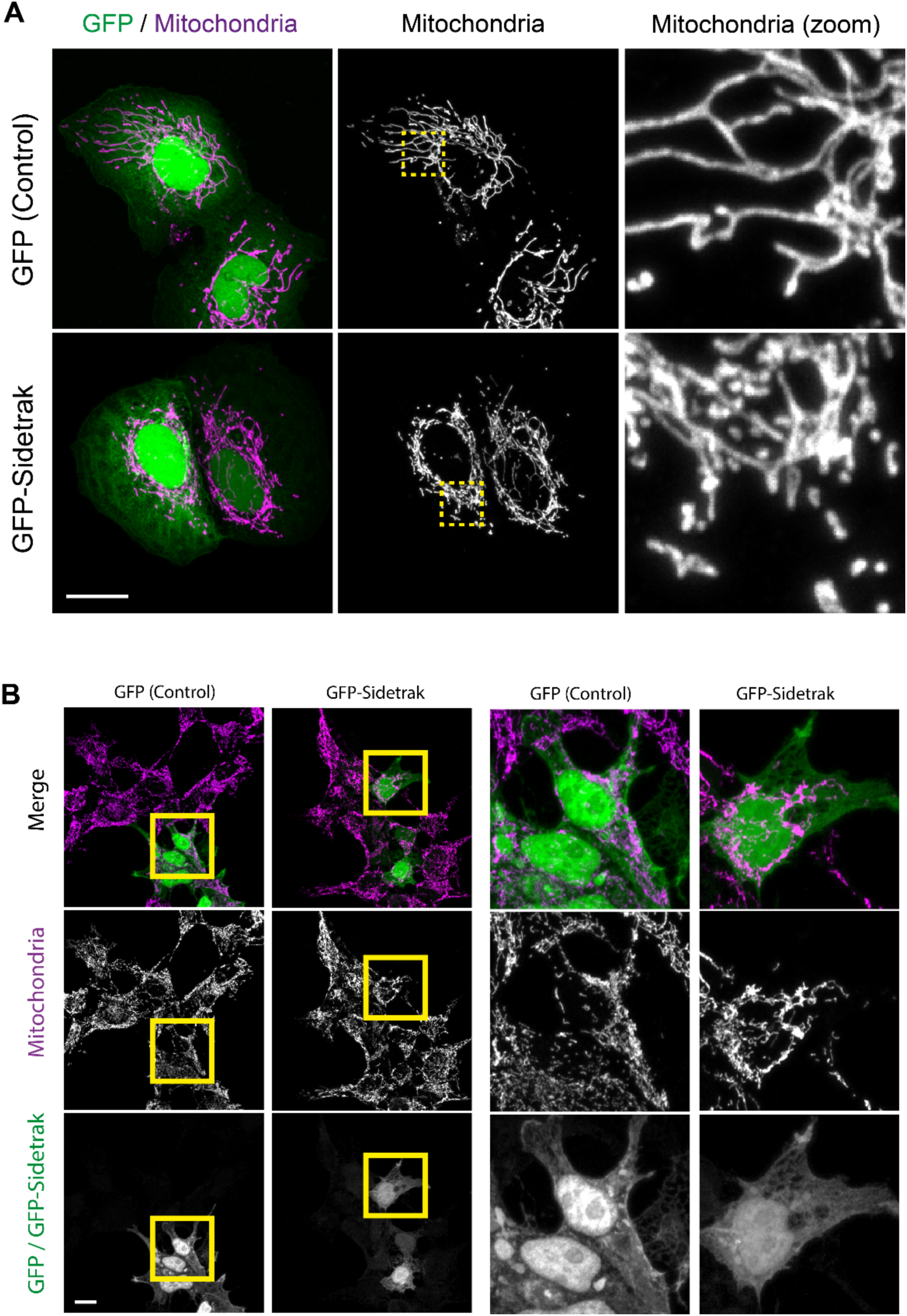
Sidetrak induces mitochondrial clustering in multiple cell types. Representative fluorescence images of (A) U2OS (B) HEK293 cells expressing GFP or GFP-Sidetrak and stained for mitochondria. Expression of GFP-Sidetrak resulted in a pronounced redistribution of mitochondria towards the perinuclear region in both cell types, similar to that observed in HeLa cells. Scale bars, 20 µm.

**Figure S12.**
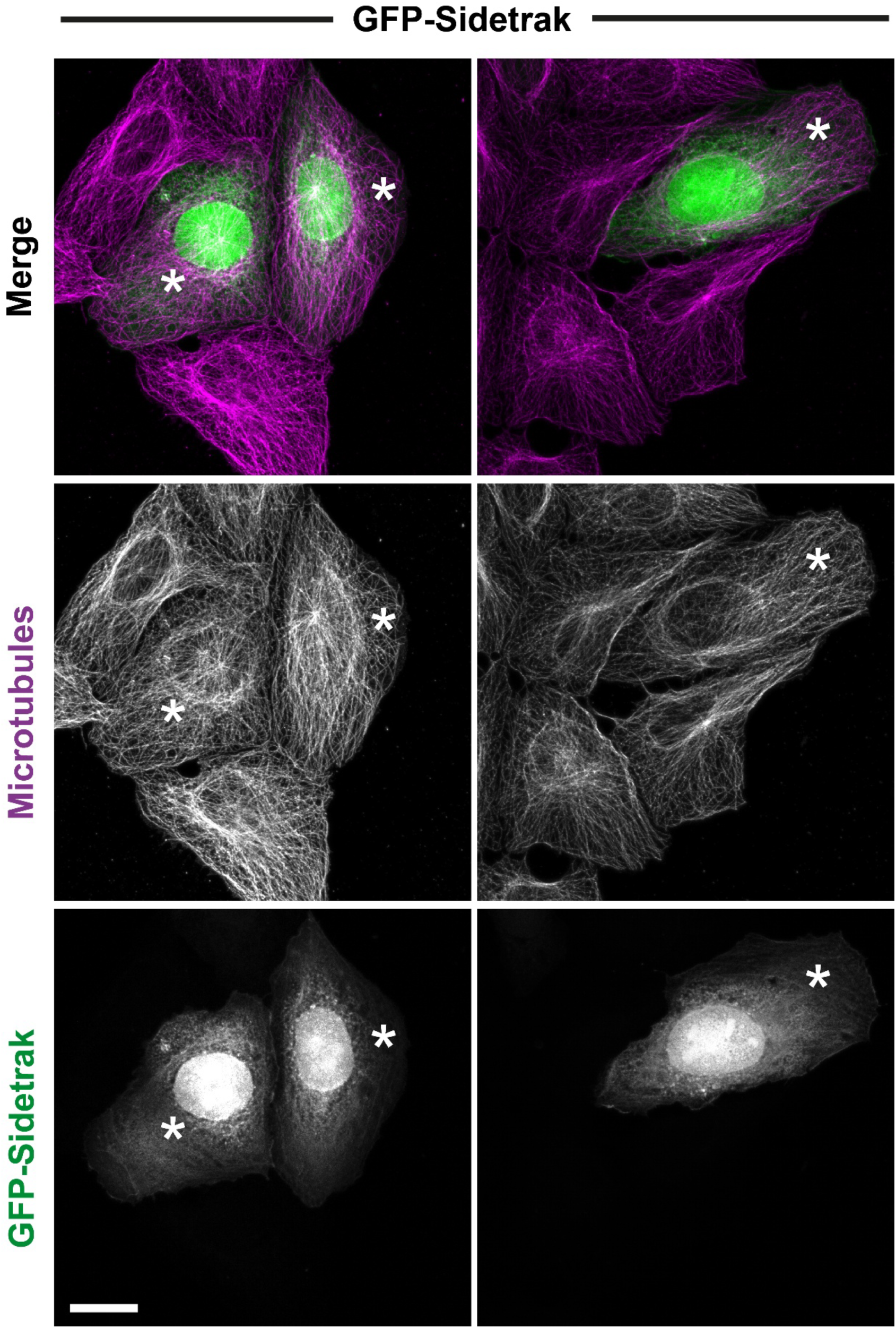
Sidetrak does not affect gross microtubule architecture. U2OS cells expressing GFP-Sidetrak were immunostained for tubulin. Cells expressing GFP-Sidetrak (*) can be compared to non-transfected cells in the same field. Scale bars, 10 µm.

**Figure S13.**
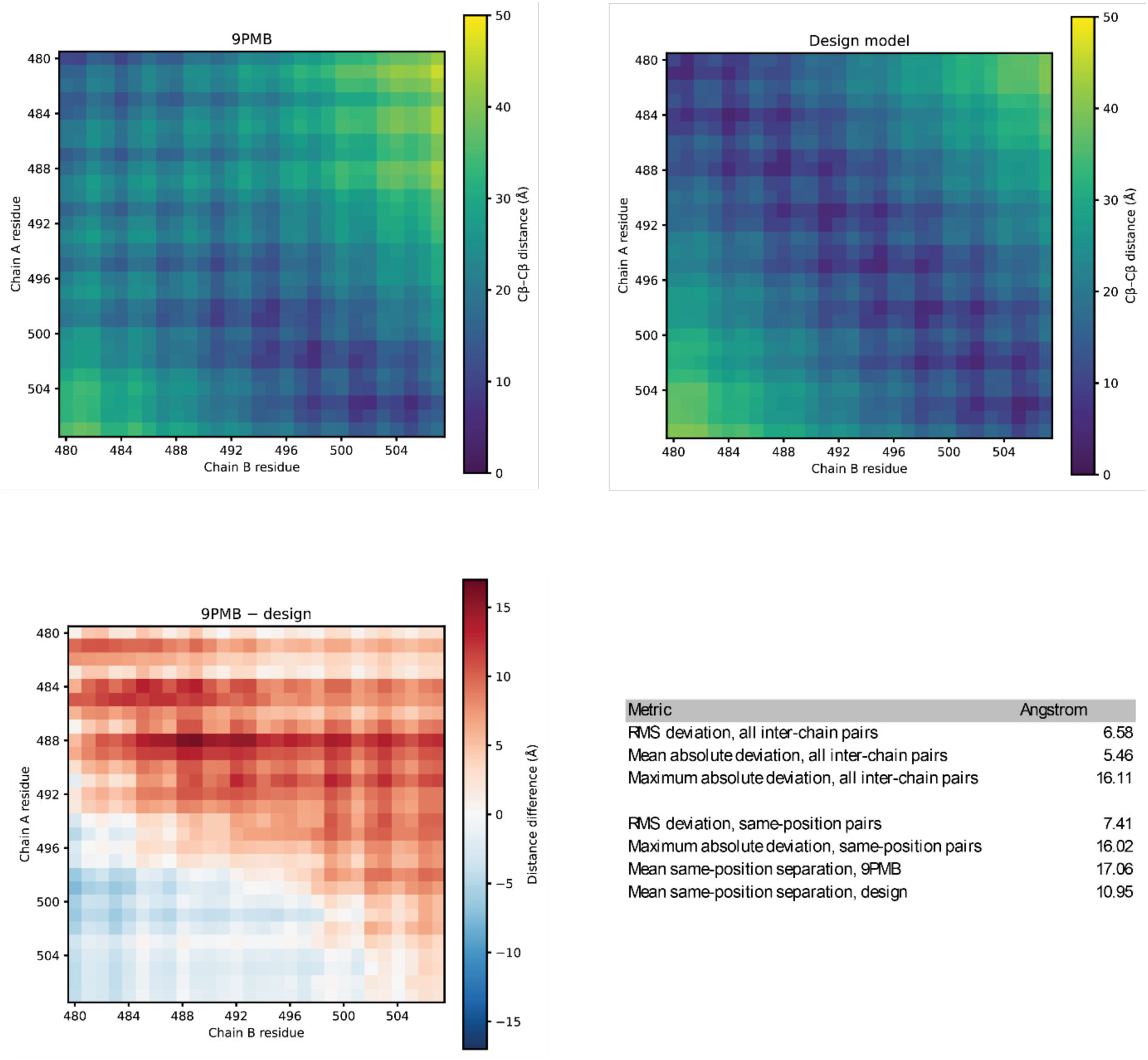
Comparison of TDS geometry in the AF3-derived design model and cryo-EM model of kinesin-1. Structural comparison of the four-heptad TDS within the AF3-derived CC1/Sidetrak design model (human KIF5B) and the deposited cryo-EM model (PDB: 9PMB). Inter-chain Cβ–Cβ distances were calculated across corresponding 28-residue regions of chains A and B and represented as pairwise distance matrices. Across all inter-chain residue pairs, the RMS difference between the two matrices was 6.58 Å, with a mean absolute difference of 5.46 Å and a maximum difference of 16.11 Å. Analysis restricted to equivalent residue positions across the two chains gave an RMS difference of 7.41 Å, with mean inter-chain separations of 10.95 Å in the AF3-derived design model and 17.06 Å in 9PMB. These analyses demonstrate substantial differences in the relative geometry of the two CC1 helices across the TDS.

## References

1. A. Yildiz, Mechanism and regulation of kinesin motors. Nat Rev Mol Cell Biol 26, 86–103 (2025).

2. S. J. Lehman, C. Crocini, L. A. Leinwand, Targeting the sarcomere in inherited cardiomyopathies. Nat Rev Cardiol **19**, 353–363 (2022).

3. K. I. Albanese, S. Barbe, S. Tagami, D. N. Woolfson, T. Schiex, Computational protein design. Nature Reviews Methods Primers 5, 13 (2025).

4. J. A. Cross, M. S. Chegkazi, R. A. Steiner, D. N. Woolfson, M. P. Dodding, Fragment-linking peptide design yields a high-affinity ligand for microtubule-based transport. Cell Chem Biol 28, 1347–1355 e1345 (2021).

5. J. A. Cross et al., A de novo designed coiled coil-based switch regulates the microtubule motor kinesin-1. Nat Chem Biol 20, 916–923 (2024).

6. J. A. Cross, D. N. Woolfson, M. P. Dodding, Controlling cell architecture with protein design. Curr Opin Cell Biol **96**, 102569 (2025).

7. J. S. Allingham, B. H. Kwok, H. Sosa, Kinesins as dynamic drug targets: structural insights and emerging therapeutic strategies. Trends in Biochemical Sciences, (2026).

8. J. A. Cross, M. P. Dodding, Motor-cargo adaptors at the organelle-cytoskeleton interface. Curr Opin Cell Biol **59**, 16–23 (2019).

9. Z. Tan, A. Missman, M. A. Cianfrocco, Rules of the road: how to turn off kinesin motors. Biochem J 483, 229–245 (2026).

10. J. F. Weijman et al., Molecular architecture of the autoinhibited kinesin-1 lambda particle. Sci Adv 8, eabp9660 (2022).

11. G. Carrington et al., A multiscale approach reveals the molecular architecture of the autoinhibited kinesin KIF5A. J Biol Chem 300, 105713 (2024).

12. S. Shukla et al., Kinesin-1 conformational dynamics are controlled by a cargo-sensitive TPR switch. Elife 14, RP109462 (2026).

13. P. J. Hooikaas et al., MAP7 family proteins regulate kinesin-1 recruitment and activation. J Cell Biol 218, 1298–1318 (2019).

14. T. Metzger et al., MAP and kinesin-dependent nuclear positioning is required for skeletal muscle function. Nature 484, 120–124 (2012).

15. K. Barlan, W. Lu, V. I. Gelfand, The microtubule-binding protein ensconsin is an essential cofactor of kinesin-1. Curr Biol 23, 317–322 (2013).

16. B. Y. Monroy et al., Competition between microtubule-associated proteins directs motor transport. Nat Commun 9, 1487 (2018).

17. S. R. Tymanskyj, B. H. Yang, K. J. Verhey, L. Ma, MAP7 regulates axon morphogenesis by recruiting kinesin-1 to microtubules and modulating organelle transport. Elife 7, e36374 (2018).

18. M. Métivier et al., Dual control of Kinesin-1 recruitment to microtubules by Ensconsin in Drosophila neuroblasts and oocytes. Development 146, (2019).

19. L. S. Ferro et al., Structural and functional insight into regulation of kinesin-1 by microtubule-associated protein MAP7. Science 375, 326–331 (2022).

20. M. Ashaduzzaman et al., Structural basis of kinesin-1 autoinhibition and its control of microtubule-based motility. Sci Adv 12, eaeg1267 (2026).

21. J. Abramson et al., Accurate structure prediction of biomolecular interactions with AlphaFold 3. Nature 630, 493–500 (2024).

22. H. Stark et al., BoltzGen: Toward Universal Binder Design. bioRxiv, 2025.2011.2020.689494 (2025).

23. A. A. Ibarra et al., Predicting and Experimentally Validating Hot-Spot Residues at Protein-Protein Interfaces. ACS Chem Biol 14, 2252–2263 (2019).

24. C. W. Wood et al., BAlaS: fast, interactive and accessible computational alanine-scanning using BudeAlaScan. Bioinformatics 36, 2917–2919 (2020).

25. K. Brickley, F. A. Stephenson, Trafficking kinesin protein (TRAK)-mediated transport of mitochondria in axons of hippocampal neurons. J Biol Chem 286, 18079–18092 (2011).

26. M. van Spronsen et al., TRAK/Milton motor-adaptor proteins steer mitochondrial trafficking to axons and dendrites. Neuron 77, 485–502 (2013).

27. K. Brickley, M. J. Smith, M. Beck, F. A. Stephenson, GRIF-1 and OIP106, members of a novel gene family of coiled-coil domain proteins: association in vivo and in vitro with kinesin. J Biol Chem 280, 14723–14732 (2005).

28. J. W. Hammond, K. Griffin, G. T. Jih, J. Stuckey, K. J. Verhey, Co-operative versus independent transport of different cargoes by Kinesin-1. Traffic 9, 725–741 (2008).

29. R. S. Stowers, L. J. Megeath, J. Gorska-Andrzejak, I. A. Meinertzhagen, T. L. Schwarz, Axonal transport of mitochondria to synapses depends on milton, a novel Drosophila protein. Neuron 36, 1063–1077 (2002).

30. A. Bracale et al., Kidins220/ARMS is transported by a kinesin-1-based mechanism likely to be involved in neuronal differentiation. Mol Biol Cell 18, 142–152 (2007).

31. X. Wang et al., KIDINS220 and InsP8 safeguard the stepwise regulation of phosphate exporter XPR1. Mol Cell 85, 3209–3224 e3208 (2025).

32. K. Brickley, K. Pozo, F. A. Stephenson, N-acetylglucosamine transferase is an integral component of a kinesin-directed mitochondrial trafficking complex. Biochim Biophys Acta 1813, 269–281 (2011).

33. S. P. Iyer, Y. Akimoto, G. W. Hart, Identification and cloning of a novel family of coiled-coil domain proteins that interact with O-GlcNAc transferase. J Biol Chem 278, 5399–5409 (2003).

34. G. Lansbergen et al., CLASPs Attach Microtubule Plus Ends to the Cell Cortex through a Complex with LL5β. Developmental Cell 11, 21–32 (2006).

35. I. Grigoriev et al., Rab6 regulates transport and targeting of exocytotic carriers. Dev Cell 13, 305–314 (2007).

36. L. Fourriere et al., RAB6 and microtubules restrict protein secretion to focal adhesions. J Cell Biol 218, 2215–2231 (2019).

37. L. Wang, K. Lee, R. Malonis, I. Sanchez, B. D. Dynlacht, Tethering of an E3 ligase by PCM1 regulates the abundance of centrosomal KIAA0586/Talpid3 and promotes ciliogenesis. Elife 5, (2016).

38. M. R. Hannaford et al., Pericentrin interacts with Kinesin-1 to drive centriole motility. J Cell Biol 221, (2022).

39. M. R. Hannaford, N. M. Rusan, Positioning centrioles and centrosomes. J Cell Biol 223, (2024).

40. M. C. Hunt, M. I. Siponen, S. E. Alexson, The emerging role of acyl-CoA thioesterases and acyltransferases in regulating peroxisomal lipid metabolism. Biochim Biophys Acta 1822, 1397–1410 (2012).

41. M. Tello-Lafoz et al., Sorting nexin 27 interactome in T-lymphocytes identifies zona occludens-2 dynamic redistribution at the immune synapse. Traffic 18, 491–504 (2017).

42. J. U. Jung, M. H. Cobb, WNK1 controls endosomal trafficking through TRIM27-dependent regulation of actin assembly. Proc Natl Acad Sci U S A 120, e2300310120 (2023).

43. N. Sahabandu et al., Active microtubule-actin cross-talk mediated by a nesprin-2G-kinesin complex. Sci Adv 11, eadq4726 (2025).

44. T. S. Randall et al., A small-molecule activator of kinesin-1 drives remodeling of the microtubule network. Proc Natl Acad Sci U S A 114, 13738–13743 (2017).

45. B. M. Bensel et al., Kinesin-1 autoinhibition tunes cargo transport by motor ensembles. Biophys J 124, 3278–3290 (2025).

46. G. G. Rhys et al., De novo designed peptides for cellular delivery and subcellular localisation. Nat Chem Biol 18, 999–1004 (2022).

47. B. Mylemans et al., De Novo-Designed Bifunctional Proteins for Targeted Protein Degradation. J Am Chem Soc 148, 34311–34319 (2026).

48. F. Kozielski et al., The crystal structure of dimeric kinesin and implications for microtubule-dependent motility. Cell 91, 985–994 (1997).

49. J. Niu et al., Kinesin-1 holoenzyme assembly coordinates cargo-adaptor recognition with heavy-chain autoinhibition. *bioRxiv*, 2026.2008.2020.746125 (2026).

50. K. Chiba, K. M. Ori-McKenney, S. Niwa, R. J. McKenney, Synergistic autoinhibition and activation mechanisms control kinesin-1 motor activity. Cell Rep 39, 110900 (2022).

51. X. Robert, C. Guillon, P. Gouet, FoldScript: a web server for the efficient analysis of AI-generated 3D protein models. Nucleic Acids Res 53, W277–W282 (2025).

52. Y. Y. Yip et al., The light chains of kinesin-1 are autoinhibited. Proc Natl Acad Sci U S A 113, 2418–2423 (2016).

53. T. D. Pollard, A guide to simple and informative binding assays. Mol Biol Cell 21, 4061–4067 (2010).

54. P. Kumar, D. N. Woolfson, Socket2: a program for locating, visualizing and analyzing coiled-coil interfaces in protein structures. Bioinformatics 37, 4575–4577 (2021).

55. A. M. Waterhouse, J. B. Procter, D. M. Martin, M. Clamp, G. J. Barton, Jalview Version 2--a multiple sequence alignment editor and analysis workbench. Bioinformatics 25, 1189-1191 (2009).

